# Gamma frequency neuronal oscillations modulate microglia morphology via colony stimulating factor 1 receptor and NFκB pathway signalling

**DOI:** 10.1101/2025.03.01.641001

**Authors:** Megan Elley, Jonathan T. Brown, Jonathan Witton

## Abstract

Within the brain, neurons and glial cells engage in dynamic crosstalk to maintain homeostasis and regulate neuroimmune responses. Recent studies have implicated rhythmic neuronal network activity, most notably at gamma oscillation frequencies (approx. 25-100 Hz), in modulating the morphology and function of microglia, the brain’s primary immune cells. Little is known, however, about the cellular mechanisms underlying this form of neuroimmune communication. Using pharmacological and optogenetic models of gamma oscillations in mouse brain slices, we found that gamma oscillations stimulate microglia to adopt a reactive morphological phenotype via activation of colony stimulation factor 1 receptor (CSF1R) and nuclear factor κB-mediated signalling. Surprisingly, inhibition of two downstream mediators of CSF1R signalling – phosphoinositide-3-kinase or phospholipase C – did not prevent this effect, suggesting that neuron-microglia interactions in this context may occur via compensatory or alternative CSF1R-linked pathways. These findings provide important insights into how rhythmic brain activity regulates neuroimmune function, with potential implications for neurological health and disease.

## Introduction

Neuronal oscillations are rhythmic electrical waves that reflect temporally structured activity in neuronal networks, emerging in response to cognitive and behavioural demands. Gamma oscillations are approximately 25-100 Hz neuronal oscillations that arise through the coordinated firing of ensembles of excitatory and fast-spiking inhibitory interneurons (1). These oscillations are observed throughout the brain and are associated with a range of higher-order cognitive and behavioural processes, including memory, emotion, sensory perception, and movement (2).

While gamma oscillations have long been recognised in neuronal communication (1), recent studies have demonstrated that they may also initiate an as-yet-uncharacterised signal that regulates the function of microglia, the resident immune cells of the central nervous system (CNS) (3–5). Microglia are highly dynamic cells that extend motile processes into the brain parenchyma to continuously survey the brain environment and respond to physiological and pathological perturbations. Upon detecting a homeostatic challenge, microglia rapidly assume an amoeboid appearance by enlarging their soma and retracting their processes to transform into a ‘reactive’ phenotype associated with phagocytosis and release of immune peptides (6). Bidirectional communication between neurons and microglia occurs via a complex network of small molecule and contact-dependent mechanisms (7), but the observation that gamma oscillations can influence neuron-microglia signalling is striking, as it suggests that the temporal organisation of neuronal network activity plays a crucial role in regulating neuroimmune function and maintaining brain homeostasis.

Studies have revealed that experimentally inducing gamma oscillations through 40 Hz optogenetic or sensory (auditory and/or visual) stimulation can drive microglial activation and reduce the abundance of pathological entities such as amyloid-β peptide (Aβ) and hyperphosphorylated tau protein in mouse models of Alzheimer’s disease (AD) (3–5). These findings led us to investigate whether this neuroimmune interaction occurs under physiological conditions and to explore the signalling mechanisms involved. Using an *ex vivo* mouse brain slice model, we demonstrate that gamma oscillations induced via established pharmacological (8) and optogenetic (9) methods stimulate microglia to adopt amoeboid morphologies characteristic of a reactive, phagocytic phenotype. Furthermore, we demonstrate that this neuroimmune interaction is mediated by activation of colony stimulating factor 1 (CSF1) receptors and the nuclear factor κ-light-chain-enhancer of activated B cells (NFκB) signalling pathway. These findings provide important insights into the mechanistic basis of gamma oscillation-driven neuron-microglia communication, with important implications for neuroimmune regulation and brain homeostasis.

## Results

### Gamma oscillations modulate microglia morphology in mouse brain slices

We investigated the effects of gamma oscillations on microglia morphology using a well-established pharmacological model of gamma oscillations in the CA3 region of hippocampus in C57BL/6J mouse cortico-hippocampal slices. As previously described (8), bath application of the muscarinic acetylcholine receptor (mAChR) agonist carbachol (CCh, 20 µM) induced gamma oscillations in the CA3 local field potential (LFP; measured at 20-50 Hz; Power: - 55.0±2.2 dB, Frequency: 31.3±4.2 Hz, N=7(4)). In contrast, gamma oscillations were absent under vehicle conditions, where CCh was not applied (CCh-condition; Power: -63.1±2.2 dB, *p*=0.020, Frequency: 20.6±0.7 Hz, *p*=0.023, N=5(2), **Table 1**; **Fig. 1A-C**). Slices were fixed after stable gamma oscillations were established (35 min post CCh) and microglia within a 100 µm radius of the LFP recording site in CA3 *stratum radiatum* were visualised using anti-Iba1 immunolabelling followed by 2-photon microscopy (**Fig. 1D**). Quantification of microglial ramification (using a Ramification Index, RI, **Methods**; **Fig. S1**) revealed that microglia exposed to CCh evoked gamma oscillations (CCh+ condition) exhibited less ramified, more amoeboid morphologies (RI: 5.84±1.36, N=70(7,4)) compared to microglia in CCh-control conditions (RI: 8.05±1.32, N=50(5,2), *p*=0.041, **Table 2**; **Fig.1E**). We replicated this experiment in heterozygous CX3CR1^+/GFP^ mice, which express enhanced green fluorescent protein (EGFP) in microglia under control of the endogenous *Cx3cr1* locus (10). Upon inducing CCh evoked gamma oscillations in CX3CR1^+/GFP^ cortico-hippocampal slices (Power: CCh+: - 57.4±2.3 dB, CCh-: -64.5±1.1 dB, *p*<0.001, Frequency: CCh+: 25.0±2.7 Hz, CCh-: 20.7±0.6 Hz, *p*<0.001, N=6(6), **Table 3**; **Fig. 1F-H**), we observed a similar reduction in the ramification of EGFP-labelled microglia at the CA3 recording site of CCh+ slices (RI: 5.00±1.03, N=60(6,6)) compared to CCh-slices (RI: 5.87±0.87, N=60(6,6), *p*<0.001, **Table 4**; **Fig. 1I,J**), confirming our finding in C57BL/6J mouse slices. As an additional control, we examined the morphology of microglia in the perirhinal cortex (PRh) of CX3CR1^+/GFP^ slices, where bath application of CCh induced a small increase in broadband (1-50 Hz) network activity, but did not evoke gamma oscillations (Power: CCh+^PRh^: -58.8±2.8 dB, CCh-^PRh^: -66.1±3.4 dB, *p*<0.001, Frequency: CCh+^PRh^: 20.7±0.8 Hz, CCh-^PRh^: 27.9±10.4 Hz, *p*=0.11, N=6(6), **Table S1**; **Fig. S2A-C**) (11). In PRh, microglia ramification did not significantly differ between CCh+ and CCh-conditions (CCh+^PRh^ RI: 6.21±1.06, N=60(6,6), CCh-^PRh^ RI: 6.46±1.18, N=60(6,6), *p*=0.74, Table S2; Fig. S2D,E).

**Figure 1.**
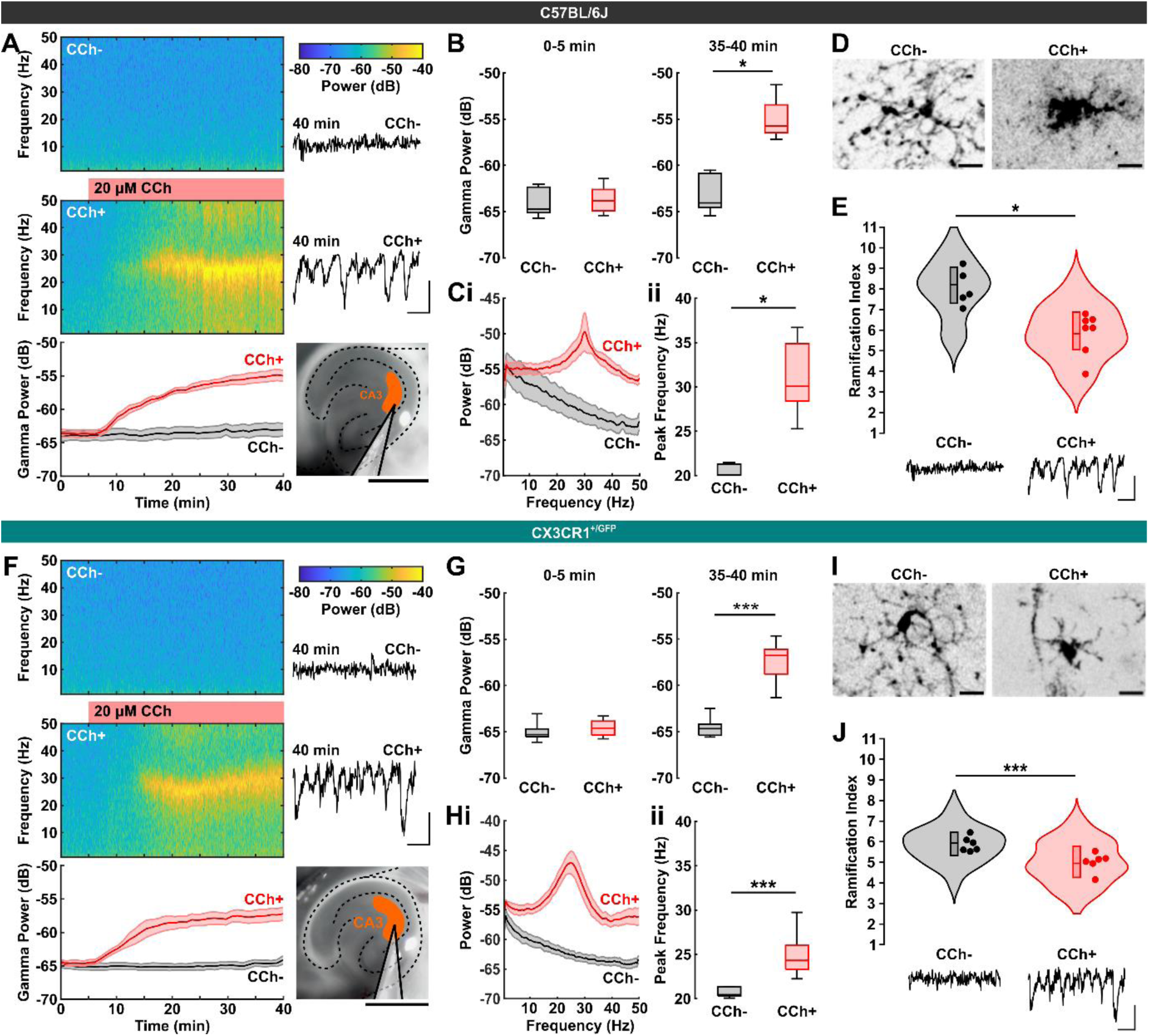
Pharmacologically evoked gamma oscillations modulate microglial morphology in mouse brain slices. **(A)** Representative spectrograms and summary time course (bottom) showing the development of gamma oscillations in CA3 of C57BL/6J slices (inset image; scale bar: 1 mm) following bath application of CCh (at 5 min; CCh+ condition; middle). Gamma oscillations were absent in slices where CCh was not applied (CCh condition; top). Inset traces illustrate LFPs at 40 min (scale bar: 0.1 mV, 50 ms). **(B)** Mean gamma band (20-50 Hz) power measured during the pre-CCh period (0-5 min) and post-CCh period (35-40 min) in C57BL/6J slices. **(C)** Power spectra (mean ± SEM) **(i)** and summary data **(ii)** showing peak gamma frequency during the post-CCh period (35-40 min) in CCh- and CCh+ C57BL/6J slices. **(D)** 2-photon micrographs of example anti-Iba1 immunolabelled microglia in the CA3 region of CCh- and CCh+ C57BL/6J slices (scale bar: 10 µm). **(E)** CA3 microglia RIs in C57BL/6J slices under CCh- and CCh+ conditions. Dots represent the mean RI per slice. Traces illustrate example LFPs per condition (scale bar: 0.1 mV, 50 ms). **(F-J)** Corresponding data for CX3CR1^+/GFP^ slices, following the same structure as panels **A-E**. In **I**, example microglia express EGFP under control of the endogenous *Cx3cr1* locus. See **Tables 1-4** for statistical analysis. \**p*<0.05, \*\*\**p*<0.001.

**Table 1.** Linear mixed effects models comparing baseline gamma power (0-5 min) and CCh evoked gamma power and frequency (35-40 min) between vehicle control (CCh-) and CCh treatment (CCh+) conditions in the CA3 region of C57BL/6J cortico-hippocampal slices. To evaluate model goodness-of-fit, the table displays Akaike Information Criterion values for null and mixed effects (condition) models. The effect of CCh treatment was evaluated by T-test. The table displays the degrees of freedom (*df*), test statistic (*t*), and significance (*p*) values. For fixed effects, the table displays the estimates and confidence intervals (CI). For random effects, the table displays residual variance (σ2), mouse variance (τ00), and the intraclass correlation coefficient (ICC). Marginal R^2^ refers to the variance explained by fixed effects only. Conditional R^2^ refers to variance explained by fixed and random effects combined.

| **Linear mixed effects model**<br>*GammaProperty ~ condition + (1 \| mouse)* |  |  |  |  |  |  |  |  |  |
| --- | --- | --- | --- | --- | --- | --- | --- | --- | --- |
|  | **Power (0-5 min)** |  |  | **Power (35-40 min)** |  |  | **Frequency (35-40 min)** |  |  |
| **Goodness-of-fit** |  |  |  |  |  |  |  |  |  |
|  | *Null* | *Condition* |  | *Null* | *Condition* |  | *Null* | *Condition* |  |
| *Akaike information criterion* | 46.36 | 48.28 |  | 55.21 | 48.16 |  | 74.55 | 67.53 |  |
| **T-Test** |  |  |  |  |  |  |  |  |  |
|  | *df* | *t* | *p* | *df* | *t* | *p* | *df* | *t* | *p* |
|  | 3.63 | 0.23 | 0.83 | 4.00 | 3.76 | **0.020** | 3.63 | 3.79 | **0.023** |
| **Fixed effects** |  |  |  |  |  |  |  |  |  |
| *Predictors* | *Estimates* | *CI* |  | *Estimates* | *CI* |  | *Estimates* | *CI* |  |
| Intercept (CCh-) | -63.73 | -66.21, 61.25 |  | -62.68 | -66.44, -58.91 |  | 20.49 | 15.35, 25.63 |  |
| CCh+ | 0.31 | -2.77, 3.39 |  | 7.54 | 2.91, 12.16 |  | 10.53 | 4.13, 16.93 |  |
| **Random effects** |  |  |  |  |  |  |  |  |  |
| $\sigma^2$ | 0.92 | | | 0.40 | | | 5.20 | | |
| $\tau_{00}$ mouse | 1.94 | | | 5.17 | | | 7.78 | | |
| Intraclass correlation coefficient | 0.68 |  |  | 0.93 |  |  | 0.60 |  |  |
| *N* mouse | 6 |  |  | 6 |  |  | 6 |  |  |
| *N* observations | 12 |  |  | 12 |  |  | 12 |  |  |
| Marginal $R^2$ | 0.009 | | | 0.730 | | | 0.694 | | |
| Conditional $R^2$ | 0.682 | | | 0.980 | | | 0.877 | | |

**Table 2.** Linear mixed effects model comparing microglia ramification indices (RI) between vehicle control (CCh-) and CCh treatment (CCh+) conditions in the CA3 region of C57BL/6J mouse cortico-hippocampal slices. To evaluate model goodness-of-fit, the table displays Akaike Information Criterion values for null and mixed effect (condition) models. The effect of CCh treatment was evaluated by T-test. The table displays the degrees of freedom (*df*), test statistic (*t*), and significance (*p*) values. For fixed effects, the table displays the estimates and confidence intervals (CI). For random effects, the table displays residual variance (σ2), mouse variance (τ00 mouse), slice variance (τ00 slice:mouse), and the intraclass correlation coefficient (ICC). Marginal R^2^ refers to the variance explained by fixed effects only. Conditional R^2^ refers to variance explained by fixed and random effects combined.

| Linear mixed effects model |  |  |  |
| --- | --- | --- | --- |
| RI ~ condition + (1 \| mouse / slice) |  |  |  |
| Goodness-of-fit |  |  |  |
|  | Null | Condition |  |
| Akaike information criterion | 392.96 | 387.62 |  |
| T-Test |  |  |  |
|  | df | t | p |
|  | 3.76 | -3.06 | 0.041 |
| Fixed effects |  |  |  |
| Predictors | Estimates | CI |  |
| Intercept (CCh-) | 8.16 | 6.93, 9.38 |  |
| CCh+ | -2.37 | -3.91, -0.83 |  |
| Random effects |  |  |  |
| $\sigma^2$ | 1.13 | | |
| $\tau_{00}$ slice:mouse | 0.39 | | |
| $\tau_{00}$ mouse | 0.56 | | |
| Intraclass correlation coefficient | 0.46 |  |  |
| N slice | 12 |  |  |
| N mouse | 6 |  |  |
| N observations | 120 |  |  |
| Marginal R<sup>2</sup> | 0.398 |  |  |
| Conditional R<sup>2</sup> | 0.674 |  |  |

**Table 3.** Linear mixed effects models comparing baseline gamma power (10-15 min) and CCh evoked gamma power and frequency (35-40 min) between vehicle control (CCh-), CCh treatment (CCh+), CCh plus atropine treatment (CCh+ATR), and CCh plus NBQX treatment (CCh+NBQX) conditions in the CA3 region of CX3CR1^+/GFP^ mouse cortico-hippocampal slices. To evaluate model goodness-of-fit, the table displays Akaike Information Criterion values for null and mixed effects (condition) models. Main fixed effects were evaluated by one-way ANOVA. The table displays the degrees of freedom (*df*), test statistics (*F*), and significance (*p*) values. For fixed effects, the table displays the estimates, confidence intervals (CI), and *p* values for *post hoc* tests (Dunnett’s comparison to CCh- with Holm-Bonferroni correction). For random effects, the table displays residual variance (σ2), mouse variance (τ00), and the intraclass correlation coefficient (ICC). Marginal R^2^ refers to the variance explained by fixed effects only. Conditional R^2^ refers to variance explained by fixed and random effects combined.

| Linear mixed effects model |  |  |  |  |  |  |  |  |  |  |  |  |
| --- | --- | --- | --- | --- | --- | --- | --- | --- | --- | --- | --- | --- |
| GammaProperty ~ condition + (1 \| mouse) |  |  |  |  |  |  |  |  |  |  |  |  |
|  | Power (10-15 min) |  |  |  | Power (35-40 min) |  |  |  | Frequency (35-40 min) |  |  |  |
| Goodness-of-fit |  |  |  |  |  |  |  |  |  |  |  |  |
|  | Null |  | Fixed Effects |  | Null |  | Fixed Effects |  | Null |  | Fixed Effects |  |
| Akaike Information Criterion | 85.00 |  | 88.57 |  | 131.25 |  | 110.96 |  | 115.60 |  | 92.44 |  |
| ANOVA |  |  |  |  |  |  |  |  |  |  |  |  |
|  | df |  | F | p | df |  | F | p | df |  | F | p |
|  | Between | 3 | 0.649 | 0.61 | Between | 3 | 17.44 | <0.001 | Between | 3 | 16.37 | <0.001 |
|  | Within | 4.97 |  |  | Within | 10.54 |  |  | Within | 19.15 |  |  |
| Fixed effects |  |  |  |  |  |  |  |  |  |  |  |  |
| Predictors | Estimates |  | CI | p | Estimates |  | CI | p | Estimates |  | CI | p |
| Intercept (CCh-) | -64.73 |  | -66.26, -63.73 |  | -64.27 |  | -65.97, -62.50 |  | 20.64 |  |  |  |
| CCh+ | 0.08 |  | -1.07, 1.07 |  | 7.01 |  | 4.88, 9.15 | <0.001 | 4.35 |  | 2.66, 6.03 | <0.001 |
| CCh+ATR | 0.91 |  | -1.31, 1.34 |  | 1.51 |  | -0.89, 3.91 | 0.42 | -0.39 |  | -2.10, 1.32 | 0.93 |
| CCh+NBQX | 0.01 |  | -0.56, 2.38 |  | 1.66 |  | -0.57, 3.89 | 0.29 | -0.40 |  | -2.09, 1.30 | 0.93 |
| Random effects |  |  |  |  |  |  |  |  |  |  |  |  |
| $\sigma^2$ | 0.79 | | | | 2.47 | | | | 1.86 | | | |
| $\tau_{00}$ mouse | 1.18 | | | | 2.24 | | | | 0.15 | | | |
| Intraclass correlation coefficient | 0.60 |  |  |  | 0.48 |  |  |  | 0.07 |  |  |  |
| N mouse | 17 |  |  |  | 17 |  |  |  | 17 |  |  |  |
| N observations | 24 |  |  |  | 24 |  |  |  | 24 |  |  |  |
| Marginal R<sup>2</sup> | 0.72 |  |  |  | 0.610 |  |  |  | 0.675 |  |  |  |
| Conditional R<sup>2</sup> | 0.627 |  |  |  | 0.796 |  |  |  | 0.699 |  |  |  |

**Table 4.**
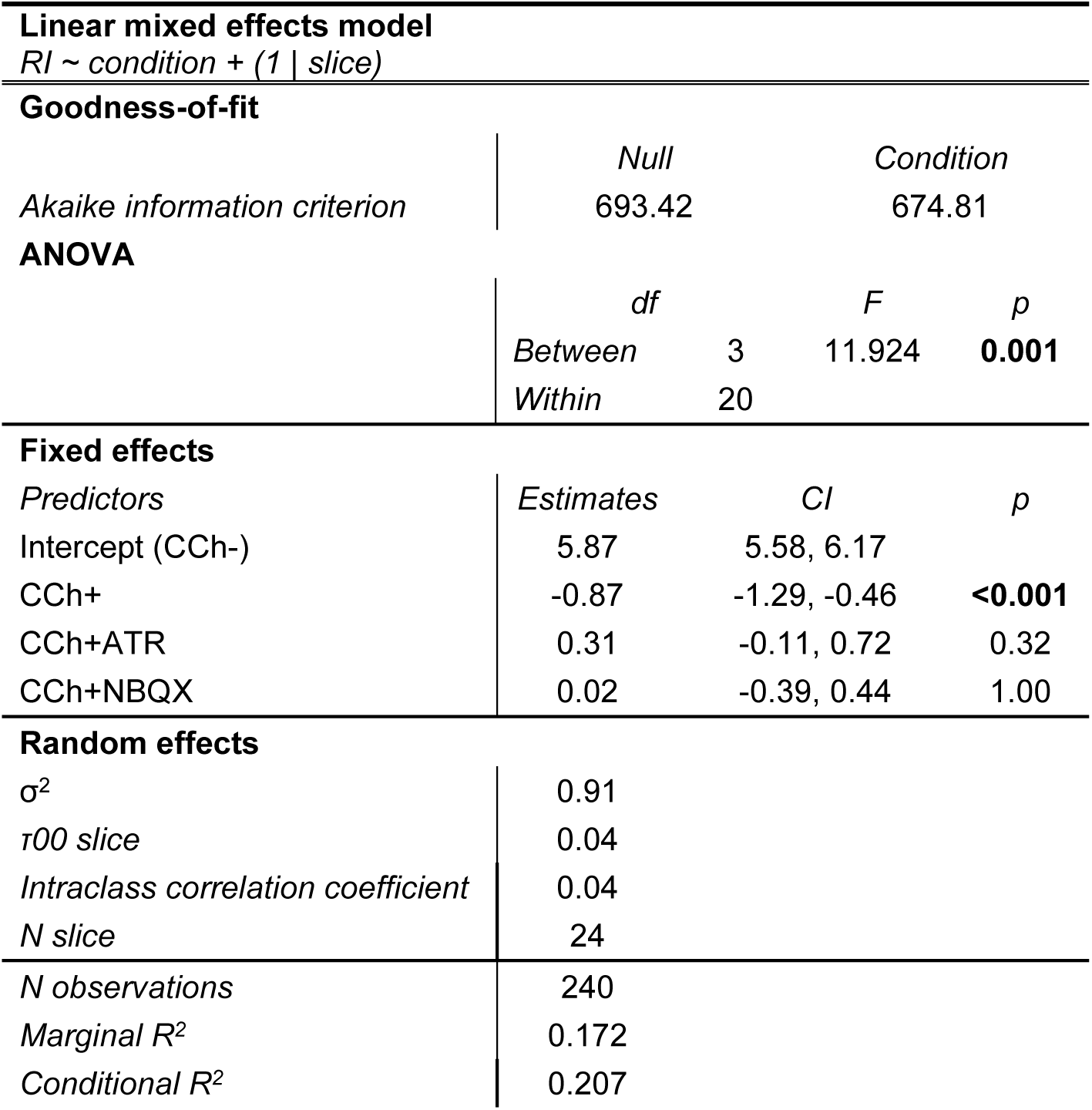
Linear mixed effects model comparing microglia ramification indices (RI) between vehicle control (CCh-), CCh treatment (CCh+), CCh plus atropine treatment (CCh+ATR), and CCh plus NBQX treatment (CCh+NBQX) conditions in the CA3 region of CX3CR1^+/GFP^ mouse cortico-hippocampal slices. To evaluate model goodness-of-fit, the table displays Akaike information criterion (AIC) values for null and mixed effects (condition) models. Main fixed effects were evaluated by one-way ANOVA. The table displays the degrees of freedom (*df*), test statistic (*F*), and significance (*p*) values. For fixed effects, the table displays the estimates, confidence intervals (CI), and *p* values for *post hoc* tests (Dunnett’s comparison to CCh- with Holm-Bonferroni correction). For random effects, the table displays residual variance (σ2), slice variance (τ00 slice), and the intraclass correlation coefficient (ICC). Marginal R^2^ refers to the variance explained by fixed effects only. Conditional R^2^ refers to variance explained by fixed and random effects combined.

Because a subpopulation (11-16%) of adult microglia express M3 muscarinic acetylcholine receptors (mAChRs) (12), we confirmed that the observed changes in microglia morphology were specifically driven by CCh evoked gamma oscillations. First, CCh was co-applied to CX3CR1^+/GFP^ slices with the non-selective mAChR antagonist atropine (10 µm; CCh+ATR condition), which effectively suppressed the induction of gamma oscillations (Power: -62.8±2.3 dB, *p*=0.42, Frequency: 20.3±0.5 Hz, *p*=0.93, N=6(6), **Table 3**; **Fig. 2A-C**). Next, we co-applied CCh with the AMPA/kainate receptor antagonist NBQX (1 µm; CCh+NBQX condition), which also prevented the induction of CCh evoked gamma oscillations without blocking mAChRs (Power: -62.3±2.5 dB, *p*=0.29, Frequency: 20.3±0.4 Hz, *p*=0.93, N=6(6), **Table 3**; **Fig. 2A-C**). Microglia in both CCh+ATR (RI: 6.18±0.88, N=60(6,6), *p*=0.32) and CCh+NBQX (RI: 5.90±1.08, N=60(6,6), *p*=1.00) conditions remained ramified, exhibiting morphologies consistent with microglia in CCh-slices (**Table 4**; **Fig. 2D,E**). Additionally, in PRh, where CCh treatment failed to induce gamma oscillations (**Table S1**; **Fig. S2A-C**), microglia morphology was comparable across CCh-^PRh^, CCh+ATR^PRh^ (RI: 6.27±0.84, N=60(6,6)), and CCh+NBQX^PRh^ (RI: 6.17±0.83, N=60(6,6)) conditions (*p*=0.74; **Table S2**; **Fig. S2D,E**).

**Figure 2.**
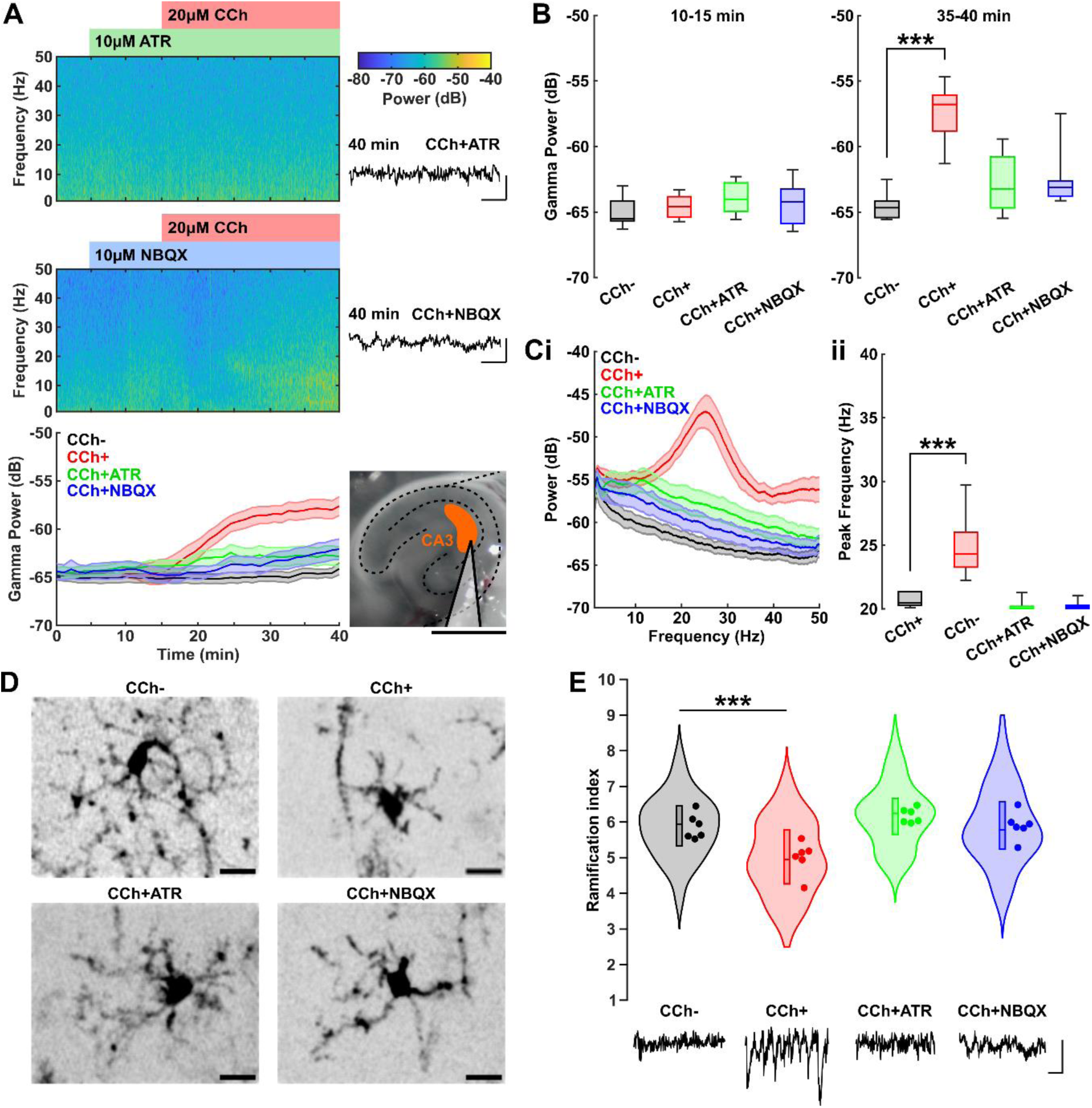
Suppression of pharmacologically evoked gamma oscillations prevents microglial morphology changes in the presence of CCh. **(A)** Representative spectrograms and summary time course (mean ± SEM; bottom) showing that atropine (ATR, top) and NBQX (middle; each applied at 5 min) supress the induction of gamma oscillations by CCh (applied at 15 min) in CA3 (inset image; scale bar: 1 mm). Inset traces illustrate LFPs at 40 min (scale bar: 0.1 mV, 50 ms). **(B)** Mean gamma band (20-50 Hz) power measured during the pre-CCh period (10-15 min) and post-CCh period (35-40 min). **(C)** Power spectra (mean ± SEM) **(i)** and summary data **(ii)** showing peak gamma frequency during the post-CCh period (35-40 min). **(D)** 2-photon micrographs of example CA3 microglia in CCh-, CCh+, CCh+ATR, and CCh+NBQX slices (scale bar: 10 µm). **(E)** CA3 microglia RIs under CCh-, CCh+, CCh+ATR, and CCh+NBQX conditions. Dots represent the mean RI per slice. Traces illustrate example LFPs per condition (scale bar: 0.1 mV, 50 ms). Treatment conditions were interleaved between brain slices. Data for the CCh- and CCh+ groups in panels **B-E** are reproduced from **Fig. 1F-J** to facilitate direct comparison with the CCh+ATR and CCh+NBQX conditions. See **Tables 3 & 4** for statistical analysis. \*\*\**p*<0.001.

In behaving animals, hippocampal gamma oscillations typically occur nested within lower frequency theta (4-12 Hz) oscillations (13). To further evaluate the effects of gamma oscillations on microglia morphology, we implemented a model of theta-coupled gamma oscillations in mouse brain slices using optogenetic stimulation (9). CX3CR1^+/GFP^ mice received injections of retrogradely transported adeno-associated virus (AAV2/retro) into the CA1 region of the hippocampus to express channelrhodopsin-2 (ChR2; H134R variant (14)) in CA3 pyramidal neurons, without activating CA3-resident microglia through AAV expression (**Fig. S3**; **Methods**). Four weeks post injection, cortico-hippocampal slices were prepared and theta (5 Hz) modulated blue (470 nm) light stimulation was used to evoke theta-coupled gamma oscillations in CA3 for 30 min (Wavelet power: 0.0314±0.0160, N=5(4), *p*=0.025, **Table 5**; **Fig. 3A-G**). Consistent with our pharmacological gamma oscillation model (**Fig. 1**), analysis of microglia morphology revealed that exposure to theta-coupled gamma oscillations resulted in less ramified microglia (RI: 5.33±0.95, N=50(5,4)) compared to control slices (RI: 6.05±1.13, N=40(4,3), p=0.016, **Table 6**; **Fig. 3H,I**), where red (625 nm) light stimulation of the same intensity (5 mW) did not evoke multiplexed theta-gamma oscillations (Wavelet power: 0.0055±0.0014, N=4(3), **Table 5**).

**Figure 3.**
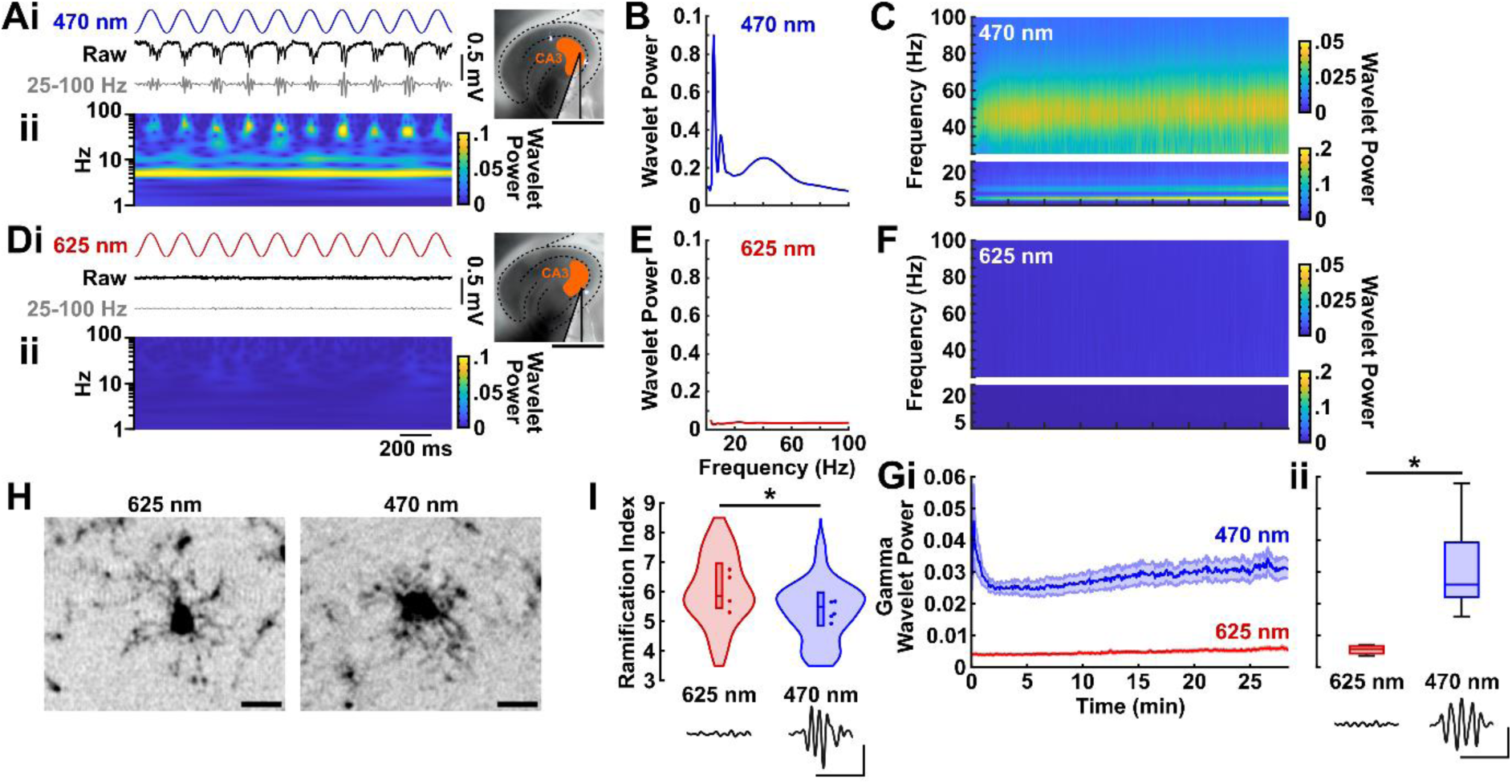
Optogenetically evoked theta-coupled gamma oscillations modulate microglial morphology in mouse brain slices. **(A)** Example traces **(i)** and continuous wavelet transform **(ii)** illustrating optogenetic induction of theta-coupled gamma oscillations in CA3 of a ChR2-transduced brain slice (inset image; scale bar: 1 mm). The traces display the theta (5 Hz) modulated 470 nm light stimulus (top), raw LFP (middle), and gamma band (25-100 Hz) filtered LFP (bottom) (scale bar: 0.1 mV; 200 ms). **(B)** Wavelet power spectrum of the LFP trace in **A**. **(C)** Wavelet spectrogram corresponding to the experiment shown in **A & B**. **(D-F)** Corresponding data for a ChR2-transduced slice stimulated with 625 nm light, following the same structure as panels **A-C**. **(G)** Time course (mean ± SEM) **(i)** and summary data **(ii)** showing that 5 Hz modulated 470 nm light stimulation induced robust theta-coupled gamma oscillations in ChR2-transduced slices, whereas 625 nm control light stimulation did not. **(H)** 2-photon micrographs of example CA3 microglia in ChR2-transduced slices exposed to 30 min theta-modulated gamma oscillations (470 nm light stimulation) and control conditions (625 nm light stimulation) (scale bar: 10 µm). **(I)** CA3 microglia RIs for theta-modulated gamma (470 nm) and control (625 nm) stimulation conditions. Dots represent the mean RI per slice. Traces in **Gii** and **I** illustrate example gamma filtered CA3 LFPs per condition (scale bar: 0.2 mV, 100 ms). See **Tables 5 & 6** for statistical analysis. \**p*<0.05.

**Table 5.** Linear mixed effects model comparing gamma wavelet power between 625 nm (control) and 470 nm light stimulation conditions in the CA3 region of ChR2-transduced CX3CR1^+/GFP^ mouse cortico-hippocampal slices. To evaluate model goodness-of-fit, the table displays Akaike Information Criterion values for null and mixed effects (condition) models. The effect of 470 nm stimulation was evaluated by T-test. The table displays the degrees of freedom (*df*), test statistic (*t*), and significance (*p*) values. For fixed effects, the table displays the estimates and confidence intervals (CI). For random effects, the table displays residual variance (σ2), mouse variance (τ00), and the intraclass correlation coefficient (ICC). Marginal R^2^ refers to the variance explained by fixed effects only. Conditional R^2^ refers to variance explained by fixed and random effects combined.

| Linear mixed effects model |  |  |  |
| --- | --- | --- | --- |
| GammaWaveletPower ~ condition + (1 \| mouse) |  |  |  |
| Goodness-of-fit |  |  |  |
|  | Null | Condition |  |
| Akaike Information Criterion | -43.10 | -48.19 |  |
| T-test |  |  |  |
|  | df | t | p |
|  | 6.15 | 2.93 | **0.025** |
| Fixed effects |  |  |  |
| Predictors | Estimates | CI |  |
| Intercept (625 nm) | 0.007 | -0.006, 0.021 |  |
| 470 nm | 0.023 | 0.007, 0.041 |  |
| Random effects |  |  |  |
| $\sigma^2$ | 0.0001 | | |
| $\tau_{00}$ mouse | 0.00004 | | |
| Intraclass correlation coefficient | 0.27 |  |  |
| N mouse | 5 |  |  |
| N observations | 9 |  |  |
| Marginal $R^2$ | 0.487 | | |
| Conditional $R^2$ | 0.627 | | |

**Table 6.** Linear mixed effects model comparing microglia ramification indices (RI) between 625 nm (control) and 470 nm light stimulation conditions in the CA3 region of ChR2-transduced CX3CR1^+/GFP^ mouse cortico-hippocampal slices. To evaluate model goodness-of-fit, the table displays Akaike Information Criterion values for null and mixed effects (condition) models. Main fixed effects were evaluated by T-test. The table displays the degrees of freedom (*df*), test statistic (*t*), and significance (*p*) values. For fixed effects, the table displays the estimates and confidence intervals (CI). For random effects, the table displays residual variance (σ2), slice variance (τ00 slice), and the intraclass correlation coefficient (ICC). Marginal R^2^ refers to the variance explained by fixed effects only. Conditional R^2^ refers to variance explained by fixed and random effects combined.

| Linear mixed effects model |  |  |  |
| --- | --- | --- | --- |
| RI ~ condition + (1 \| mouse) |  |  |  |
| Goodness-of-fit |  |  |  |
|  | Null | Fixed effects |  |
| Akaike Information Criterion | 266.99 | 262.99 |  |
| T test |  |  |  |
|  | df | t | p |
|  | 57.74 | -2.48 | 0.016 |
| Fixed effects |  |  |  |
| Predictors | Estimates | CI |  |
| Intercept (625 nm) | 6.06 | 5.54, 6.59 |  |
| 470 nm | -0.63 | 1.14, -0.13 |  |
| Random effects |  |  |  |
| $\sigma^2$ | 0.94 | | |
| $\tau_{00}$ mouse | 0.18 | | |
| Intraclass correlation coefficient | 0.16 |  |  |
| N mouse | 5 |  |  |
| N slice | 9 |  |  |
| N observations | 90 |  |  |
| Marginal R<sup>2</sup> | 0.082 |  |  |
| Conditional R<sup>2</sup> | 0.231 |  |  |

These results demonstrate that experimentally induced gamma oscillations promote changes in microglia morphology, driving them to adopt a less ramified phenotype in mouse brain slices.

### Gamma oscillations modulate microglia morphology via CSF1R and NFκB signalling

Colony Stimulating Factor 1 receptors (CSF1Rs) are primarily expressed by microglia in the CNS, where they play crucial roles in microglia survival and activation states (15). Transcriptomic analysis has shown that one hour of 40 Hz *in vivo* optogenetic stimulation induces a several-fold increase in *Csf1* and *Csf1r* gene expression in mice (3), while one hour of 40 Hz flickering visual stimulation elevates CSF1 protein levels in mouse visual cortex (16). These findings suggest that CSF1Rs may be a key mediator of gamma oscillation-induced neuron-microglia signalling. To investigate this, we examined whether CCh evoked gamma oscillations could modulate microglia morphology when CSF1R signalling was suppressed using the selective inhibitors BLZ-945 (0.5 µm) (17) and pexidartinib (1 µm) (14), respectively.

We first confirmed that BLZ-945 (RI: 7.04±0.91, *p=*0.92, N=70(7,7)) and pexidartinib (RI: 7.03±0.94, *p=*0.96, N=50(5,5)) did not independently modulate microglia morphology (**Table S3**; **Fig. S4**). We then co-applied BLZ-945 (CCh+BLZ condition) or pexidartinib (CCh+PEX condition) during the induction of CCh evoked gamma oscillations in the CA3 region of CX3CR1^+/GFP^ slices and evaluated microglia ramification within 100 µm of the LFP recording site. Notably, neither BLZ-945 nor pexidartinib altered CCh evoked gamma oscillations, as both the power and frequency of gamma oscillations was consistent between CCh+ (Power: - 54.71±3.5 dB, Frequency: 26.84±3.52 Hz, N = 13(13)) and CCh+BLZ slices (Power: - 53.61±4.73 dB, *p*=0.97, Frequency: 26.24±2.35 Hz, *p*=0.69, N=7(7)), and between CCh+ and CCh+PEX slices (Power: -50.91±4.25 dB, *p*=0.19, Frequency: 23.97±3.52 Hz, *p*=0.09, N=5(5); **Table 7**; **Fig. 4A-C**). However, compared to microglia in the CCh+ condition (RI: 5.25±1.16, N=130(13,13)), microglia in CCh+BLZ slices (RI: 6.27±1.18, N=70(7,7), *p*=0.02) and CCh+PEX slices (RI: 6.87±0.84, N=50(5,5), *p*<0.001) exhibited a significantly more ramified phenotype, similar to that observed in CCh-conditions where gamma oscillations were absent (RI: 6.73±1.08, N=60(6,6), **Table 8**; **Fig. 4D,E**).

**Figure 4.**
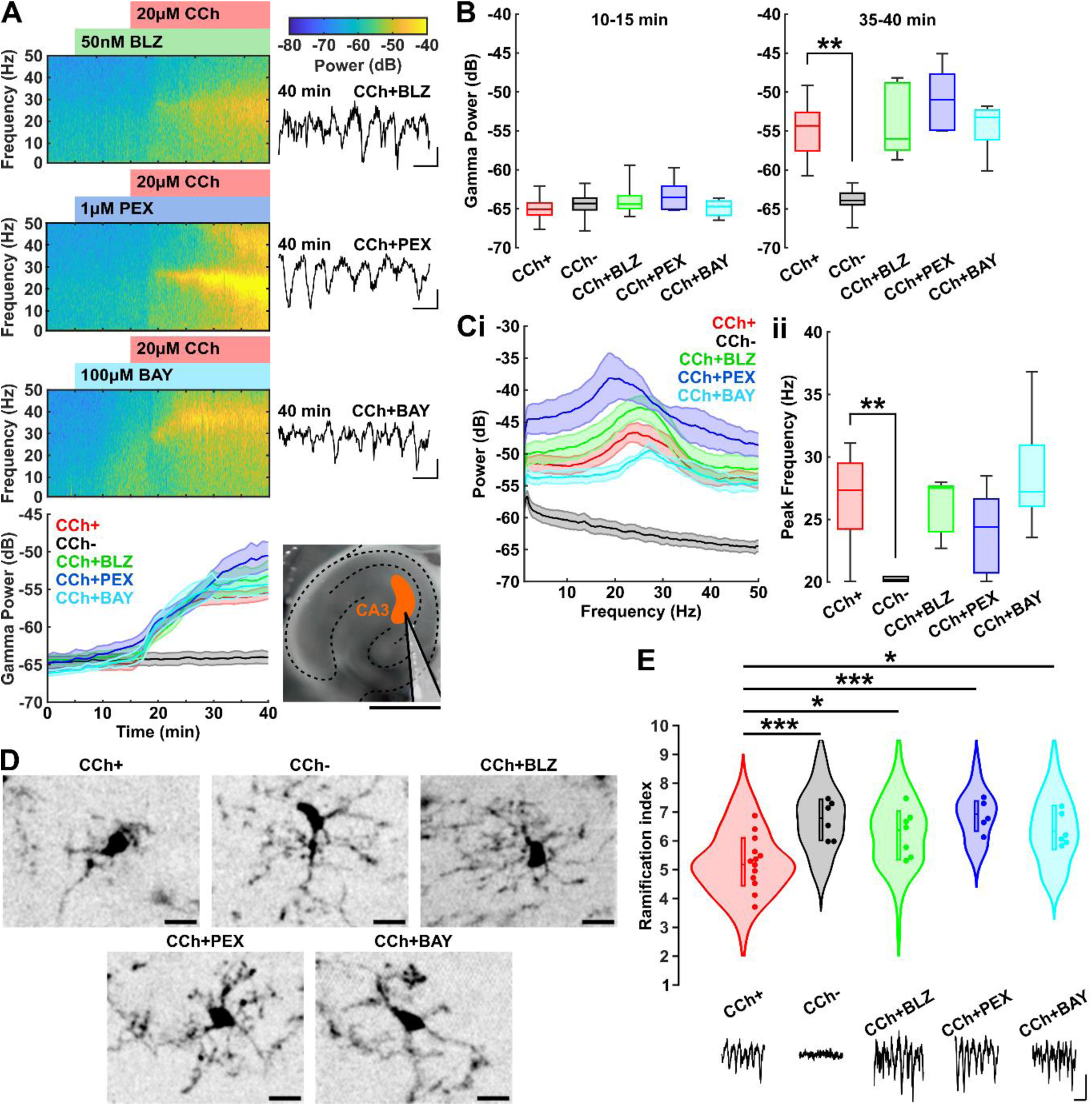
Inhibition of CSF1R or NFκB signalling supresses the modulation of microglial morphology induced by CCh evoked gamma oscillations. **(A)** Representative spectrograms and summary time course (bottom) showing that inhibition of CSF1Rs with BLZ-945 (BLZ; top) or pexidartinib (PEX; upper middle), and inhibition of the NFκB pathway with the IκK inhibitor bay 11-7082 (BAY; lower middle) (each applied at 5 min) does not disrupt gamma oscillations induced by CCh (applied at 15 min) in CA3 (inset image; scale bar: 1 mm). Inset traces illustrate LFPs at 40 min (scale bar: 0.1 mV, 50 ms). **(B)** Mean gamma band (20-50 Hz) power measured during the pre-CCh period (10-15 min) and post-CCh period (35-40 min). **(C)** Power spectra (mean ± SEM) **(i)** and summary data **(ii)** showing peak gamma frequency during the post-CCh period (35-40 min). **(D)** 2-photon micrographs of example CA3 microglia in CCh+, CCh-, CCh+BLZ, CCh+PEX and CCh+BAY slices (scale bar: 10 µm). **(E)** CA3 microglia RIs under CCh+, CCh-, CCh+BLZ, CCh+PEX, and CCh+BAY conditions. Dots represent the mean RI per slice. Traces illustrate example LFPs per condition (scale bar: 0.1 mV, 50 ms). Treatment conditions were interleaved between brain slices. See **Tables 7 & 8** for statistical analysis. *p<0.05, **p<0.01, ***p<0.001.

**Table 7.**
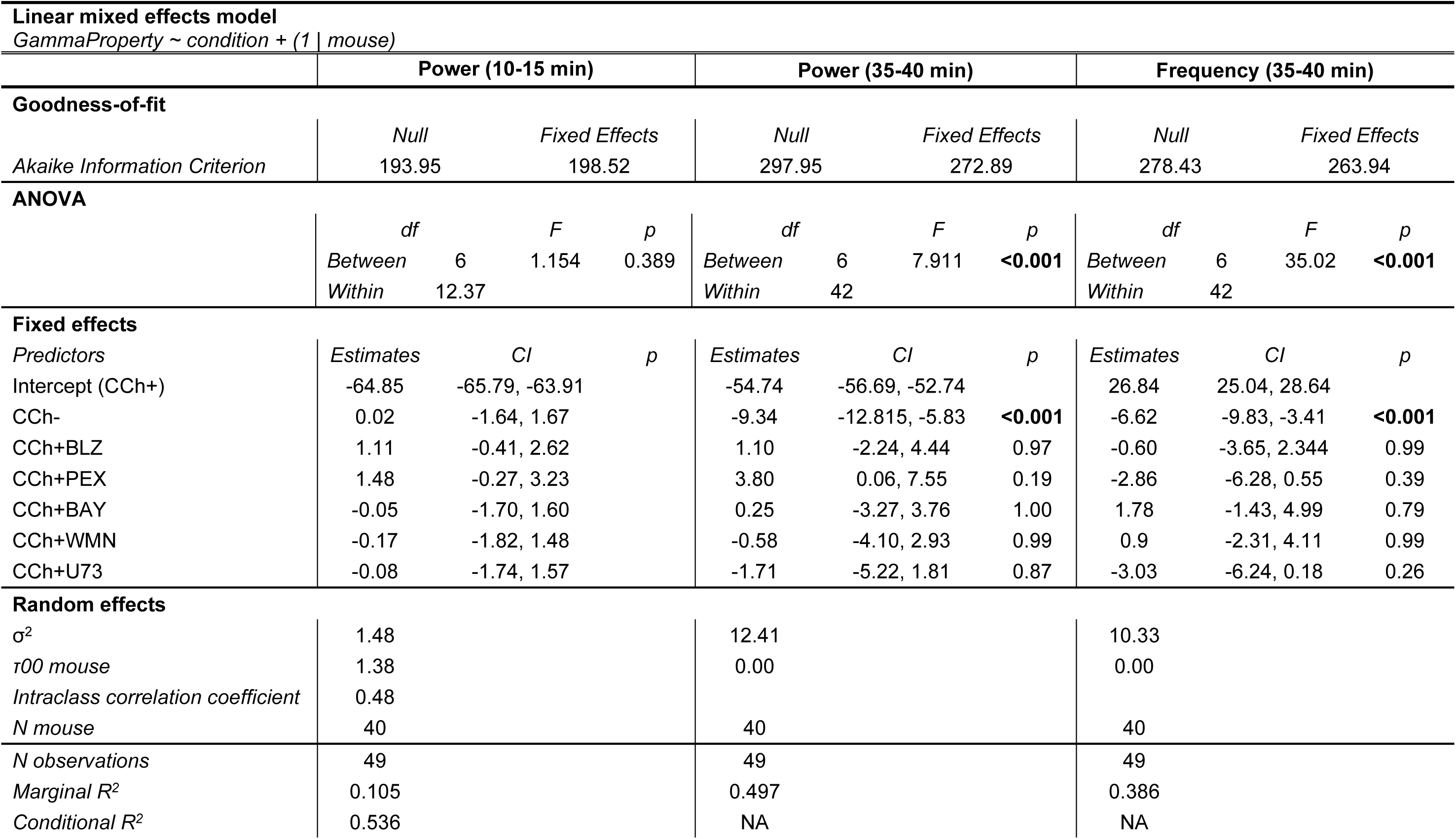
Linear mixed effects models comparing baseline gamma power (10-15 min) and CCh evoked gamma power and frequency (35-40 min) between CCh treatment (CCh+), vehicle control (CCh-), CCh plus BLZ-945 treatment (CCh+BLZ), CCh plus pexidartinib treatment (CCh+PEX), CCh plus bay 11-7082 treatment (CCh+BAY), CCh plus wortmannin treatment (CCh+WMN), and CCh plus U73122 treatment (CCh+U73) conditions in the CA3 region of CX3CR1^+/GFP^ mouse cortico-hippocampal slices. To evaluate model goodness-of-fit, the table displays Akaike Information Criterion values for null and mixed effects (condition) models. Main fixed effects were evaluated by one-way ANOVA. The table displays the degrees of freedom (*df*), test statistic (*F*), and significance (*p*) values. For fixed effects, the table displays the estimates, confidence intervals (CI), and *p* values for *post hoc* tests (Dunnett’s comparison to CCh+ with Holm-Bonferroni correction). For random effects, the table displays residual variance (σ2), mouse variance (τ00), and the intraclass correlation coefficient (ICC). Marginal R^2^ refers to the variance explained by fixed effects only. Conditional R^2^ refers to variance explained by fixed and random effects combined.

**Table 8.** Linear mixed effects model comparing microglia ramification indices (RI) between CCh treatment (CCh+), vehicle control (CCh-), CCh plus BLZ-945 treatment (CCh+BLZ), CCh plus pexidartinib treatment (CCh+PEX), CCh plus bay 11-7082 treatment (CCh+BAY), CCh plus wortmannin treatment (CCh+WMN), and CCh plus U73122 treatment (CCh+U73) conditions in the CA3 region of CX3CR1^+/GFP^ mouse cortico-hippocampal slices. To evaluate model goodness-of-fit, the table displays Akaike Information Criterion values for null and mixed effects (condition) models. Main fixed effects were evaluated by one-way ANOVA. The table displays the degrees of freedom (*df*), test statistic (*F*), and significance (*p*) values. For fixed effects, the table displays the estimates, confidence intervals (CI), and *p* values for *post hoc* tests (Dunnett’s comparison to CCh+ with Holm-Bonferroni correction). For random effects, the table displays residual variance (σ2), slice variance (τ00 slice), and the intraclass correlation coefficient (ICC). Marginal R^2^ refers to the variance explained by fixed effects only. Conditional R^2^ refers to variance explained by fixed and random effects combined.

| Linear mixed effects model |  |  |  |  |
| --- | --- | --- | --- | --- |
| Variable ~ condition + (1 \| slice) |  |  |  |  |
| Goodness-of-fit |  |  |  |  |
|  | Null |  | Fixed effects |  |
| Akaike Information Criterion | 1398.6 |  | 1379.7 |  |
| ANOVA |  |  |  |  |
|  | df |  | F | p |
|  | Between | 6 | 5.62 | <0.001 |
|  | Within | 42 |  |  |
| Fixed effects |  |  |  |  |
| Predictors | Estimates (RI) | CI (RI) | p |  |
| Intercept (CCh+) | 5.26 | 4.85 – 5.66 |  |  |
| CCh- | 1.48 | 0.75 – 2.20 | <0.001 |  |
| CCh+BLZ | 1.02 | 0.33 – 1.71 | 0.0202 |  |
| CCh+PEX | 1.62 | 0.84 – 2.39 | <0.001 |  |
| CCh+BAY | 1.11 | 0.38 – 1.83 | 0.0151 |  |
| CCh+WMN | 0.01 | -0.71 - 0.74 | 1.00 |  |
| CCh+U73 | 0.62 | -0.11 - 1.34 | 0.405 |  |
| Random effects |  |  |  |  |
| σ² | 0.79 |  |  |  |
| τ00 slice | 0.48 |  |  |  |
| Intraclass correlation coefficient | 0.38 |  |  |  |
| N slice | 49 |  |  |  |
| N observations | 490 |  |  |  |
| Marginal R² | 0.233 |  |  |  |
| Conditional R² | 0.523 |  |  |  |

NFκB proteins are a family of eukaryotic immune-regulatory transcription factors that can be activated downstream of CSF1R signalling (18). In addition to increasing levels of specific cytokines, such as CSF1, 40 Hz flickering visual stimulation has been shown to enhance phosphorylation of NFκB proteins in mice (16), suggesting that NFκB signalling may contribute to gamma oscillation-induced changes in microglial function. NFκB activation is regulated by the phosphorylation of its inhibitory binding protein, IκB, through the action of IκB kinase (IκK) (19). We therefore tested whether NFκB signalling modulates microglia morphology in our CX3CR1^+/GFP^ slice model by inducing CCh evoked gamma oscillations while co-applying the IκK inhibitor bay 11-7082 (100 µm; CCh+BAY condition) (20). Control experiments confirmed that bay 11-7082 did not independently modify microglia morphology (RI: 6.39±0.95, *p*=0.71, N=60(6,6), **Table S3**; **Fig. S4**) or alter the power (-54.47±3.22 dB, *p*=1.00) or frequency (28.62±4.68 Hz, *p*=0.26, N=6(6)) of CCh evoked gamma oscillations (**Table 7**; **Fig. 4A-C**). However, similar to the effects observed with BLZ-945 and pexidartinib, microglia in the CCh+BAY condition were significantly more ramified (RI: 6.36±1.12, N=60(6,6), *p*=0.01) than microglia exposed to gamma oscillations in the CCh+ condition, exhibiting morphologies similar to microglia in untreated CCh-slices (**Table 8**; **Fig. 4D,E**).

These findings indicate that gamma oscillations regulate microglia morphology through CSF1R and NFκB dependent signalling.

### PI3K and PLC do not mediate gamma oscillation-induced neuron-microglia signalling downstream of CSF1R activation

The CSF1R is a receptor tyrosine kinase that activates multiple molecular pathways, including those that interact with IκK/NFκB (21). In macrophages, CSF1R signalling is also known to couple with the phagocytic cell surface receptor complex formed by Triggering Receptor Expressed on Myeloid cells 2 (TREM2) and DNAX-Activating Protein of 12 kDa (DAP12) (22). Specifically, CSF1R activation triggers Src kinase-mediated phosphorylation of DAP12 when TREM2 is opsonin-bound, leading to the recruitment of downstream effectors such as phosphoinositide-3-kinase (PI3K) and phospholipase C (PLC), which promote phagocytosis of TREM2-bound substances and enhance NFκB signalling (23). Since the TREM2/DAP12 complex facilitates microglial phagocytosis of Aβ (24), and gamma frequency neuronal stimulation (3) and CSF1 administration (25) have been shown to promote Aβ clearance in mice, we hypothesised that PI3K and/or PLC may mediate gamma oscillation-induced neuron-microglia signalling downstream of CSF1Rs.

To test this, we used our CX3CR1^+/GFP^ slice model to induce CCh evoked gamma oscillations in the CA3 region of hippocampus while co-applying the PI3K inhibitor wortmannin (10 µM; CCh+WMN condition) (26) or the PLC inhibitor U73122 (10 µM; CCh+U73 condition) (27). Neither wortmannin (RI: 6.08±0.97, *p*=0.12, N=60(6,6)) nor U73122 (RI: 6.50±0.79, *p*=0.91, N=60(6,6)) independently altered microglia morphology (**Table S3**; **Fig. S4**), nor did they affect gamma oscillation properties (CCh+WMN, Power: -55.29±3.42, *p*=0.99, Frequency: 27.74±3.73, *p*=0.57, N=6(6); CCh+U73, Power: -56.42±2.71, *p*=0.87, Frequency: 23.81±2.11, *p*=0.06, N=6(6), **Table 7**; **Fig. 5A-C**). Morphological analysis of CA3 microglia revealed that cells in the CCh+WMN condition (RI: 5.27±1.15, N=60(6,6), *p*=1.00) and the CCh+U73 condition (RI: 5.87±0.93, N=60(6,6), *p*=0.40) closely resembled those in the CCh+ group, maintaining a more amoeboid appearance than microglia in CCh-slices that were not exposed to gamma oscillations (**Table 8**; **Fig. 5D,E**). Thus, inhibition of PI3K or PLC did not block gamma oscillation-induced changes in microglia morphology.

**Figure 5.**
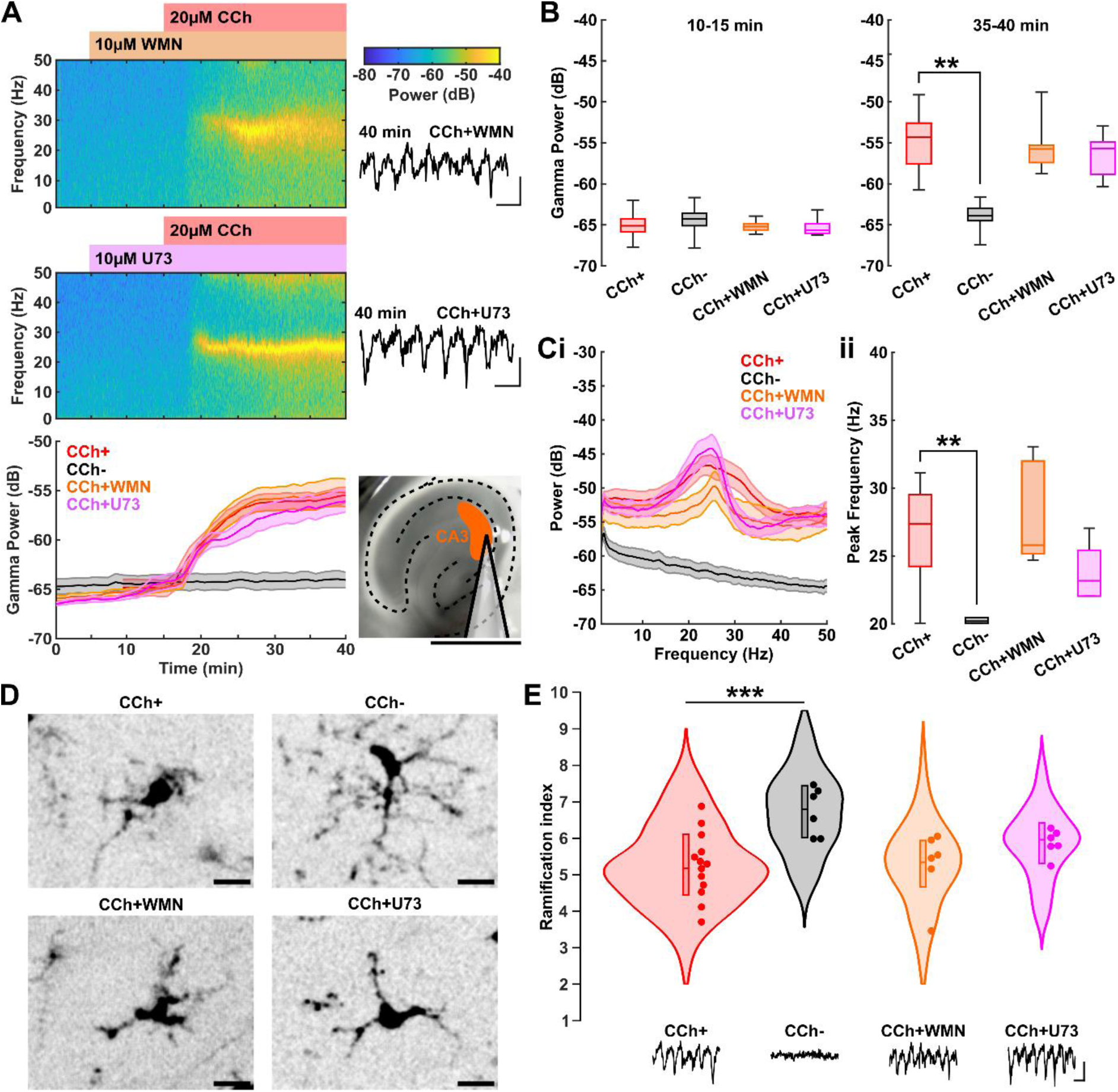
Inhibition of PI3K or PLC signalling does not affect the modulation of microglial morphology induced by CCh evoked gamma oscillations. **(A)** Representative spectrograms and summary time course (bottom) showing that inhibition of PI3K with wortmannin (WMN; top) or inhibition of PLC with U73122 (U73; middle) (each applied at 5 min) does not disrupt gamma oscillations induced by CCh (applied at 15 min) in CA3 (inset image; scale bar: 1 mm). Inset traces illustrate LFPs at 40 min (scale bar: 0.1 mV, 50 ms). **(B)** Mean gamma band (20-50 Hz) power measured during the pre-CCh period (10-15 min) and post-CCh period (35-40 min). **(C)** Power spectra (mean ± SEM) **(i)** and summary data **(ii)** showing peak gamma frequency during the post-CCh period (35-40 min). **(D)** 2-photon micrographs of example microglia in CCh+, CCh-, CCh+WMN, and CCh+U73 slices (scale bar: 10 µm). **(E)** CA3 microglia RIs under CCh+, CCh-, CCh+WMN, and CCh+U73 conditions. Dots represent the mean RI per slice. Traces illustrate example LFPs per condition (scale bar: 0.1 mV, 50 ms). Treatment conditions were interleaved between brain slices. Data for the CCh+ and CCh-groups in panels **B-E** are reproduced from Fig. 4B**-E** to facilitate direct comparison with the CCh+WMN and CCh+U73 conditions. See **Tables 7 & 8** for statistical analysis. \*\**p*<0.01, \*\*\**p*<0.001.

## Discussion

In this study, we developed a rodent brain slice assay to investigate cellular mechanisms underlying Gamma Activity-Induced Neuron-microglia Signalling (hereinafter referred to as GAINS). Our findings demonstrate that gamma oscillations modulate microglia by stimulating these cells to adopt more amoeboid morphologies, and that this process requires activation of the CSF1R and NFκB signalling pathway. Notably, we also found that PI3K and PLC signalling pathways are not transducers of microglial CSF1R activation in this context.

GAINS has previously been investigated using *in vivo* mouse models, where gamma oscillations were experimentally induced using 40 Hz optogenetic or sensory (visual and/or auditory) stimulation (3,4). Here, we investigated this phenomenon using complementary pharmacological (8) and optogenetic (9) models of gamma oscillations in mouse cortico-hippocampal brain slices. This *ex vivo* approach provided an experimentally tractable system for dissecting GAINS mechanisms using pharmacological tools. We found that microglia exhibited a less ramified, more amoeboid morphology in regions of slices where gamma oscillations were evoked (in the CA3 region of hippocampus), but not where gamma oscillations were not evoked (in PRh). These findings align with previous *in vivo* studies reporting 40 Hz optogenetic or sensory stimulation drives morphological changes in microglia, including increased soma size and reduced process length and branching (3,4). Thus, our results suggest that GAINS mechanisms are conserved between *in vivo* to *ex vivo* preparations, although further research is needed to establish whether microglia modulated by GAINS in our *ex vivo* assay exhibit comparable transcriptomic and functional profiles to those observed *in vivo* (3,16).

Previous studies investigating GAINS have sought to identify specific frequencies of neuronal oscillations that modulate microglia. These studies have indicated that GAINS may be specifically mediated by 40 Hz gamma oscillations, as neither higher frequency 80 Hz gamma oscillations nor 20 Hz beta frequency oscillations induced changes in microglia morphology (4,28). While compatible with these findings, our results suggest that GAINS may extend across a broader range of frequencies within the *low gamma* frequency band (13), as we observed a qualitatively similar modulation of microglia morphology in response to pharmacologically evoked gamma oscillations with peak frequency around 25-30 Hz.

Our study provides strong evidence that gamma oscillations signal to microglia via CSF1Rs. CSF1Rs are almost exclusively expressed by microglia in the mature CNS (14,29), and gamma oscillations failed to alter microglia morphology in the presence of the CSF1R inhibitors BLZ-945 and pexidartinib (**Fig. 4**). This finding complements prior indirect evidence implicating CSF1R signalling in GAINS. For example, one hour of 40 Hz optogenetic stimulation has been shown to upregulate *Csf1* and *Csf1r* gene expression by approximately 4-fold and 3-fold, respectively (3), while one hour of 40 Hz flickering visual stimulation led to a several-fold increase in CSF1 protein levels in the mouse visual cortex (16). Collectively, these findings suggest that CSF1R activation – likely via the cytokine CSF1 – is a key signalling mechanism linking gamma oscillations to microglia modulation. In the brain, CSF1 is produced by neurons, astrocytes, oligodendrocytes, and microglia (30). However, evidence suggests that microglia are unlikely to be a primary source of CSF1 in GAINS, as 40 Hz flickering visual stimulation has been shown to elevate CSF1 levels in both wildtype and microglia-depleted mice (28). Furthermore, astrocytes have been identified as a principal source of CSF1 in the forebrain (31), leading us to speculate that gamma oscillations may promote CSF1 release via indirect signalling through astrocytes. While further research is needed to substantiate this claim, it is noteworthy that sensory stimulation evoked gamma oscillations modulate astrocytic Ca^2+^ signalling (32), possibly through volume transmission (33).

In addition to CSF1, CSF1Rs are activated by the cytokine interleukin-34 (IL-34), which, despite lacking sequence homology with CSF1, shares a similar tertiary structure and binds to a distinct but overlapping site on CSF1Rs (34). IL-34 is predominantly released by neurons in the CNS and is highly expressed throughout the forebrain in mature animals, with limited spatial overlap with CSF1 (except in CA3 of hippocampus) (35). IL-34 and CSF1 activate similar but not identical CSF1R-mediated signalling cascades in macrophages, and several proteins generated by IL-34/CSF1R signalling – such as the chemokine eotaxin/CCL11 and IL-10 (36,37) – have been shown to be upregulated following one hour of 40 Hz flickering visual stimulation (16,28). Additionally, there is substantial overlap between genes modulated by one hour of *in vivo* 40 Hz optogenetic stimulation and those regulated by IL-34 in microglia (3). Notably, IL-34 has also been shown to enhance microglial clearance of oligomeric Aβ in neuron-microglia co-cultures (38). Thus, CSF1R activation in GAINS may also be mediated by gamma oscillation driven IL-34 release from neurons.

Beyond the role of CSF1Rs in GAINS, we found that inhibiting NFκB pathway signalling with the IκB kinase inhibitor bay 11-7082 prevented CCh evoked gamma oscillations from altering microglia morphology. This result aligns with previous studies implicating NFκB in GAINS (16,28). It has recently been proposed that gamma oscillations may selectively activate NFκB pathway signalling in neurons, as 40 Hz flickering visual stimulation has been shown to increase phosphorylated NFκB levels in neurons but not in microglia, alongside upregulation of NFκB pathway-related genes in neurons (28). Our findings are compatible with this hypothesis. However, because bay 11-7082 induced pan-cellular inhibition of NFκB in our assay – and given that NFκB is one of several pathways activated by CSF1R signalling – our findings do not rule out the possibility that gamma oscillations may also trigger NFκB activation in microglia downstream of CSF1R activation.

To investigate GAINS mechanisms resulting from CSF1R activation, we examined the effects of inhibiting key mediators of canonical CSF1R phospho-signalling pathways – specifically, PI3K (a critical component of the PI3K-Akt pathway) and PLC. These targets were selected due to their established roles in microglial phagocytosis (including Aβ clearance via interactions with TREM2/DAP12), as well as their signalling overlap with NFκB (21). Surprisingly, we found that inhibiting PI3K or PLC did not prevent gamma oscillation-induced changes in microglia ramification, suggesting that this effect is mediated by alternative CSF1R-linked pathways, or that interactions between molecular cascades downstream of CSF1Rs can compensate for selective inhibition of PI3K or PLC. A possible alternative pathway is the mitogen-activated protein kinase (MAPK) pathway, another key signalling cascade activated downstream of CSF1Rs. Recent studies suggest that MAPK phospho-signalling may play a role in GAINS (16,28). However, cell-type specific transcriptomic and gene ontology analyses indicate that 40 Hz visual stimulation upregulates MAPK pathway signalling in neurons but not in microglia (28). Moreover, MAPK pathway inhibition failed to block changes in microglia ramification induced by 40 Hz visual stimulation (28), suggesting that MAPK may be involved in GAINS, but that its effects are likely neuron-specific. Further research is therefore needed to identify molecular mediators of GAINS downstream of microglial CSF1Rs.

Several studies have reported 40 Hz optogenetic or sensory stimulation can reduce Aβ and tau pathologies and/or enhance synaptic and cognitive function in mouse models of AD (3– 5,39–41). As a result, harnessing GAINS by entraining neuronal activity at 40 Hz has been proposed as a potential therapeutic approach for AD, with upregulation of microglial phagocytosis posited as a mechanism of pathology reduction (42). While we did not directly assess microglia function, our findings are generally supportive of this hypothesis, as we demonstrate that gamma oscillations induce microglia to adopt more amoeboid morphologies characteristic of a phagocytic phenotype (43). However, this does not exclude other mechanisms by which 40 Hz stimulation may reduce protein neuropathologies, such as suppression of β-sectrease-1 cleavage of amyloid precursor protein (41). Additionally, and importantly, these effects have not been universally replicated. For instance, contrary to prior reports (4,5), Soula et al. found that that 40 Hz flickering visual stimulation failed to entrain gamma oscillations beyond the visual cortex, did not modulate microglia properties, nor decrease Aβ burden in the 5XFAD model of AD (44). Similarly, Yang and Lai (2023) reported no significant effect of 40 Hz flickering visual stimulation on microglia density or Aβ deposition in 5XFAD mice (45). These discrepancies highlight the need for further research to clarify the reliability and therapeutic potential of GAINS-based interventions.

## Concluding remarks

Our research provides valuable mechanistic insights into how gamma oscillations regulate microglia properties. We show that gamma oscillations, via CSF1R- and NFκB-dependent signalling, drive microglia in mouse cortico-hippocampal brain slices to adopt amoeboid morphologies characteristic of an ‘activated’, phagocytic phenotype. Understanding this novel form of neuron-microglia communication could open new avenues for treating neurological diseases linked to dysregulated neuroimmune function.

## Acknowledgements

We thank Aimee Sharkey, Rachel Brewer, and Matt Isherwood for their expert technical support. We also thank Dr Giselle Cheung, Dr Thomas Piers, and Prof Wendy Noble for helpful comments on the manuscript.

## Author contributions

The project was conceived and designed by ME, JTB, and JW. Experiments and analyses were performed by ME. Mouse lines were bred by ME and JW. JTB and JW supervised the work and assisted experiments and analyses carried out by ME. The manuscript was drafted by JW and ME with discussions and revisions from JTB.

## Data availability

The data and analysis code are available from the corresponding authors upon reasonable request.

## Methods

### Ethical approval

All procedures were carried out in accordance with the UK Animals (Scientific Procedures) Act 1986 and were approved by the University of Exeter Animal Welfare and Ethical Review Body.

### Animals

C57BL/6J, CX3CR1^GFP/GFP^, and CX3CR1^+/GFP^ mice (10) were bred at the University of Exeter. Homozygous CX3CR1^GFP/GFP^ and heterozygous CX3CR1^+/GFP^ mice were bred on a C57BL/6J background. Mice were group housed under standardised conditions (22±2 °C; 45±15 % humidity) and a 12-hour light/dark cycle with *ad libitum* food and water. 8-20-week-old male (M) and female (F) C57BL/6J (N=2M, 3F) and CX3CR1^+/GFP^ (N=56M, 48F) mice were used for experiments.

### Surgical procedures

For experiments described in **Fig. 3** and **Tables 5 & 6**, CX3CR1^+/GFP^ mice underwent aseptic surgery to express ChR2 (H134R variant) (46) in the hippocampus. Mice were anaesthetised with isoflurane (4-5% induction, 1-2.5% maintenance) in oxygen, and carprofen (5 mg/kg) was administered preoperatively for analgesia. Mice were secured in a stereotaxic frame, and the body temperature was maintained at ∼37 °C using a homeothermic blanket. The skull was exposed, and a craniotomy was drilled at coordinates (in mm, from Bregma): -1.65 (anteroposterior); +1.2 (mediolateral). A 33-gauge Hamilton needle was lowered into the brain to 1.2 mm below the pial surface, and 0.25-0.5 µl of AAV2/retro-CaMKII-ChR2(H134R)-mCherry (University of Zurich Vector Core) was infused into the CA1 region of the hippocampus at 0.1 µl/min. 5 min post infusion, the needle was slowly withdrawn, and the skin sutured. Following recovery from anaesthesia, mice were returned to the home cage and received postoperative carprofen (5 mg/kg in strawberry jelly) for 5 days.

### Induction and recording of gamma oscillations

Mice were euthanised by cervical dislocation, and their brains were rapidly extracted and placed in ice cold carbogenated (95% O_2_, 5% CO_2_) cutting solution containing (in mM): 189 sucrose, 10 glucose, 26 NaHCO_3_, 3 KCl, 5 MgSO_4_.7H_2_O, 0.1 CaCl_2_, 1.25 NaH_2_PO_4_. The cerebellum was removed, and 400 µm horizontal hippocampal slices were cut using a vibratome (Leica VT1200S). Slices were then hemisected and stored in an interface chamber (Digitimer), where they were continuously perfused at 30±1 °C with carbogenated artificial cerebrospinal fluid (aCSF) containing (in mM): 124 NaCl, 3 KCl, 24 NaHCO_3_, 1.25 NaH_2_PO_4_, 1 MgSO_4_.7H_2_O, 10 glucose, 2 CaCl_2_. For recordings, slices were transferred to an interface recording chamber perfused with carbogenated aCSF (2.5 ml/min) at 30±1 °C. aCSF filled glass recording electrodes (3-7 MΩ) were positioned in *stratum radiatum* of the CA3 region of the hippocampus and in layer 2/3 of PRh. Electrode placements were documented with a digital camera mounted on a stereomicroscope. LFPs were lowpass filtered at 1 kHz, amplified 200-fold using an Axopatch 200B amplifier (Molecular Devices), and digitised at 2 kHz with an analogue-to-digital converter (National Instruments). 50 Hz mains noise was removed online with a Humbug noise eliminator (Digitimer). LFPs were continuously recorded using WaveSurfer software (https://wavesurfer.janelia.org/) running in MATLAB (MathWorks).

Hippocampal gamma oscillations were induced using one of two methods. In most experiments, following a 5-15 min baseline recording, the muscarinic agonist CCh was bath-applied at 20 µM (8). For experiments performed in ChR2-transduced mice, slices were prepared 4 weeks after infusion of AAV encoding ChR2 into the hippocampus. ChR2 was activated using a 470 nm light-emitting diode (LED; Thorlabs M470F4) connected to a 400 µm diameter fibre optic patch cable positioned 2 mm above the slice using a micromanipulator. First, an input-output protocol was run to confirm that optically and electrically evoked field potentials exhibited similar amplitudes and kinetics within the same slice. Theta-coupled gamma oscillations were then elicited using a 5 Hz sinusoidal light stimulus (5 s sweeps: 4.5 s on, 0.5 s off), as previously described (9). The maximum light power at the sample was 5 mW. Control stimulation in ChR2-transduced slices was delivered at the same irradiance using a 625 nm LED (Thorlabs M625F2).

Following recordings, slices were lifted from the recording chamber on lens tissue, immediately fixed in 4% paraformaldehyde (PFA) in 0.01M phosphate buffered saline (PBS) for 1 hour, and then stored in PBS containing 0.05% sodium azide at 2.5°C.

### Pharmacology

Details of the compounds used in this study are listed in **Table S4**. All compounds were bath-applied to slices in aCSF. Different treatment conditions for the pharmacology experiments presented in **Figs. 2/S2** and in **Figs. 4/5/S5**, respectively, were interleaved between brain slices across successive recording sessions.

### Immunolabelling of microglia in C57BL/6J slices

PFA fixed, free-floating cortico-hippocampal slices were washed 6 x 5 min in Tris-buffered saline (0.025 M Tris, 0.5 M NaCl, pH 7.5) containing 0.5% TritonX-100 (TBS-T). Slices were then blocked and permeablised in 5% goat serum in TBS-T for 2 hours at room temperature (RT). Following blocking, slices were incubated overnight at RT with rabbit anti-Iba1 primary antibody (1:1000; Fujifilm) in 3% goat serum in TBS-T. Slices were then washed 6 x 5 min in TBS-T before being incubated for 4 hours at RT with Alexa Fluor 488-conjugated anti-rabbit IgG secondary antibody (1:750; Abcam) in 3% goat serum in TBS-T. Finally, slices were washed 6 x 5 min in TBS-T, mounted onto cavity microscope slides (ThermoFisher) and coverslipped using immunofluorescence mounting medium (Dako).

### Microglia imaging

Immunolabelled C57BL/6J slices and CX3CR1^+/GFP^ slices used for electrophysiological recordings were imaged using a 2-photon microscope (Scientifica) equipped with galvanometric scan mirrors and Ti:Sapphire laser (Coherent Chameleon Ultra). The laser wavelength was set to 770 nm for imaging anti-Iba1 Alexa Fluor 488-labelled microglia in C57BL/6J slices, or to 910 nm for imaging EGFP labelled microglia in CX3CR1^+/GFP^ slices. Laser power at the sample remained below 40 mW. Fluorescence was detected using a 505-545 nm emission filter and GaAsP photomultiplier tube. The microscope was controlled using ScanImage 5.0 software (Vidrio Technologies) running in MATLAB. Image stacks were acquired using a 16x objective lens (Nikon CFI Plan Fluorite, 0.8 NA) at a resolution of 512 x 512 pixels with 5x digital zoom, yielding image dimensions of 167 µm x 167 µm with a Z-step size of 0.5 µm.

### Electrophysiology data analysis

Electrophysiology data were analysed using MATLAB. LFPs were highpass filtered at 0.5 Hz and downsampled to 1 kHz. For pharmacologically induced gamma oscillations, spectrograms were generated using multi-taper spectral analysis (5 tapers, 5 s window length) via the Chronux toolbox (47). Spectral power was expressed in decibels (dB), and mean gamma oscillation power and frequency were assessed between 20-50 Hz in the 5 min preceding CCh application (baseline) and during the final 5 minutes of CCh application, once gamma oscillations had reached a steady state. For optogenetically induced theta-coupled gamma oscillations, gamma power was quantified using the continuous Morlet wavelet transform of the LFP.

### Image analysis

Images were analysed using custom routines in ImageJ (National Institutes of Health) and MATLAB. Image stacks were initially processed by subtracting the background from each Z slice using a 30 pixel rolling ball average filter. The average intensity projection of the stack was then calculated, followed by the application of a 3x3 pixel median filter to remove speckle noise. The projection image was then used to isolate regions of interest (ROIs) around the perimeter of individual microglia. 10 microglia were analysed per brain region (CA3 or PRh) per brain slice.

Microglia ramification was quantified following established methods (48). Briefly, an ROI was binarized using a manual threshold that isolated the microglia, including their processes, from the background. The perimeter and area of the cell were measured, and a ramification index (RI) was calculated as the ratio of the cell’s perimeter to its area, normalised by the same ratio for a circle of equivalent area, following

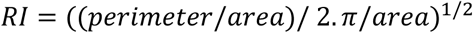

Thus, a perfect circle has an RI of 1, with higher values of RI indicating more highly ramified microglia. Representative examples of microglia with an RI between 2.9 and 8 are shown in **Fig. S1**.

### Statistics

Microglia are a heterogenous cell type, with their morphology and physiology shaped by both their endogenous state and local environment (49,50). Consequently, a given region of brain parenchyma contains phenotypically diverse microglia, which may respond differently to changes in neuronal circuit activity, such as gamma oscillations (49). However, microglia measured within the same brain slice or animal are unlikely to be fully statistically independent. Similarly, the properties of gamma oscillations evoked in different brain slices from the same animal may not be fully independent. To control for the hierarchical structure of the data (i.e., gamma power, gamma frequency, or microglia cell nested within slice and mouse), datasets were analysed using mixed effects models that included slice and/or mouse as random factors.

Data distributions were visually inspected and tested for normality using the Shapiro-Wilk test. Linear mixed models were used to analyse the data (R, v4.2.1, lme4 package). A null random intercept model was first fit, with mouse or brain slice nested within mouse as a random factor. A mixed effects model, including fixed effects of condition, was then applied. The Akaike Information Criterion was used to assess model goodness-of-fit (51). The contribution of random factors (mouse and/or slice) to the variance was also evaluated and reported (see **Tables 1-8 & S1-S3**). Statistical significance (*p*<0.05) of main fixed effects was determined using T-tests or analysis of variance (ANOVA). Degrees of freedom were estimated using Satterthwaite approximation. Where main effects were observed, *post hoc* comparisons to a defined control group were performed using Dunnett’s test, with Holm-Bonferroni adjustment for multiple comparisons (52). These effects are reported in **Tables 1-8 & S1-S3**. Pharmacology experiments presented in **Figs. 4 & 5**, respectively, were analysed using a single ANOVA.

Unless otherwise stated, descriptive statistics are presented as mean ± standard deviation or represented in box plots displaying the median, lower and upper quartiles (box), and range (whiskers). Microglia RI distributions are shown using violin plots, with overlaid dots indicating the mean RI per slice. Gamma oscillation time series are graphed as mean ± standard error of the mean (SEM). Sample sizes (N) are reported as N_slices_(N_mice_) or N_cells_(N_slices_,N_mice_), as appropriate.

---

**Figure S1.**
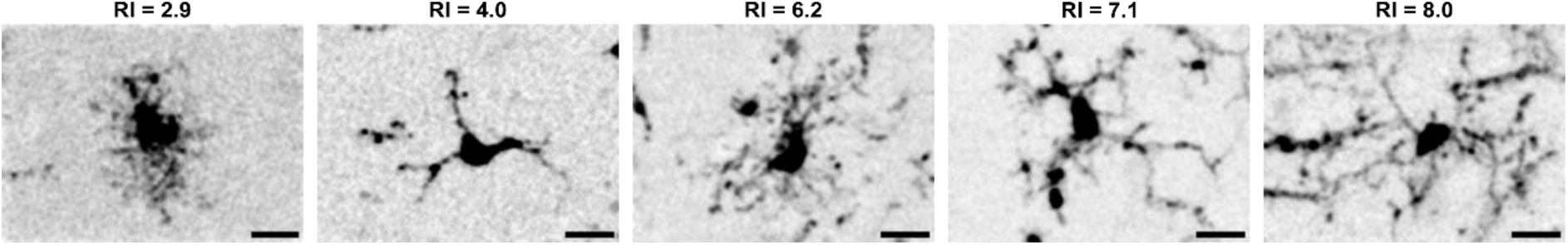
Representative CA3 microglia and corresponding ramification indices. Example 2-photon micrographs of CA3 microglia with varying ramification indices (RIs). Microglia with more amoeboid morphologies exhibit low RI values, whereas more ramified microglia display higher RIs. An RI of 1 corresponds to a perfect circle. Scale bars: 10 µm.

**Figure S2.**
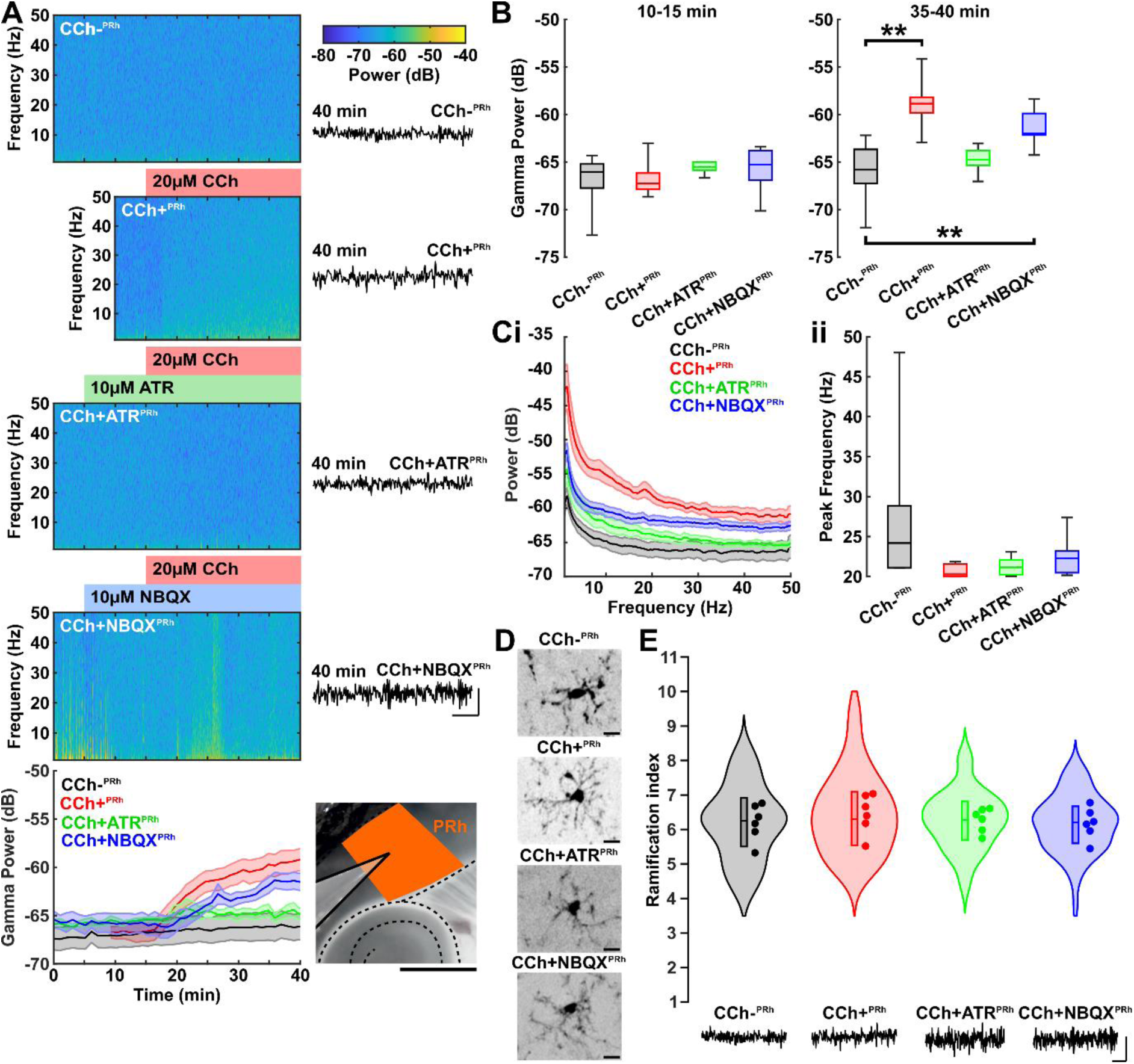
Carbachol, atropine, or NBQX do or alter microglial morphology in mouse brain slices. **(A)** Representative spectrograms and summary time course (bottom) from the PRh region of CX3CR1^+/GFP^ slices (inset image; scale bar: 1 mm), showing that bath application of carbachol (CCh+^PRh^) induces a modest increase in spectral power (+7.3 dB) compared to slices without CCh application (CCh-^PRh^), but does not generate gamma oscillations. This CCh evoked power increase is suppressed by co-application of atropine (CCh+ATR^PRh^) but is unaffected by co-application of NBQX (CCh+NBQX^PRh^) (each applied at 5 min). **(B)** Mean gamma band (20-50 Hz) power during the pre-CCh period (10-15 min) and post-CCh period (35-40 min). \*\**p*<0.01. **(C)** Power spectra (mean ± SEM) **(i)** and summary data **(ii)** showing the absence of a gamma frequency spectral peak in the post-CCh period (35-40 min) in CCh-^PRh^, CCh+^PRh^, CCh+ATR^PRh^, and CCh+NBQX^PRh^ slices. **(D)** 2-photon micrographs of example microglia in CCh-^PRh^, CCh+^PRh^, CCh+ATR^PRh^, and CCh+NBQX^PRh^ slices (scale bars: 10 µm). **(E)** Microglia RIs under CCh-^PRh^, CCh+^PRh^, CCh+ATR^PRh^, and CCh+NBQX^PRh^ conditions. Dots represent the mean RI per slice. Traces in **A & E** illustrate example LFPs per condition (scale bars: 0.1 mV, 50 ms). Treatment conditions were interleaved between brain slices. See **Tables S1 & S2** for statistical analysis.

**Figure S3.**
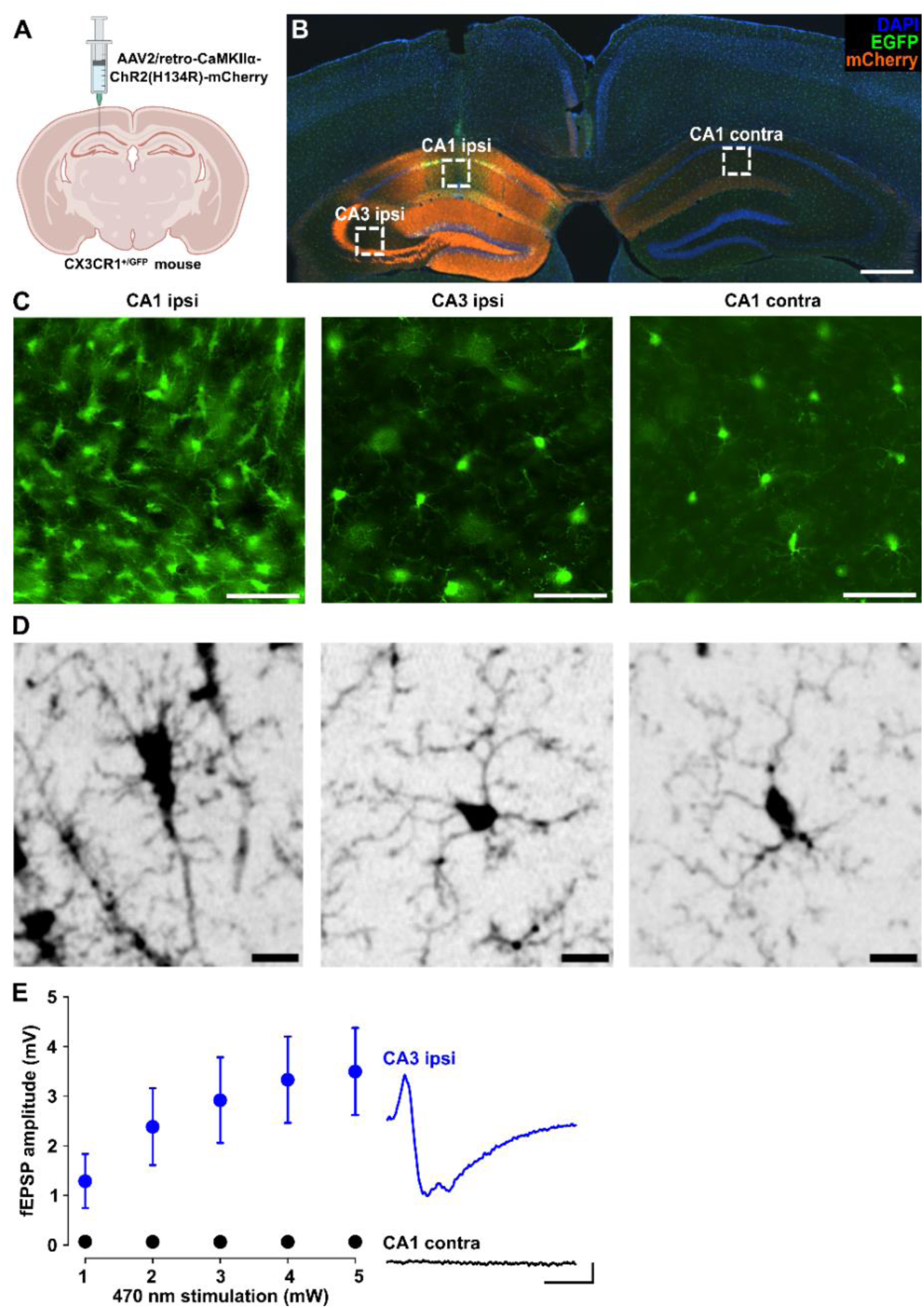
Targeted expression of ChR2 in hippocampal CA3 pyramidal neurons without AAV-induced microglia activation. **(A)** Schematic of the viral injection procedure. **(B)** Epifluorescence photomicrograph illustrating expression of AAV2/retro-CaMKIIα-ChR2(H134R)-mCherry-WPRE-hGHp(A) in the hippocampus of a CX3CR1^+/GFP^ mouse (scale bar: 500 µm). Boxes indicate regions corresponding to the CA1 injection site (CA1^ipsi^), ipsilateral CA3 expression site (CA3^ipsi^), and contralateral CA1 (CA1^contra^), which are shown at higher magnification in panels **(C)** and **(D)**. **(C)** Epifluorescence photomicrographs showing increased microglia density at the CA1 injection site (CA1^ipsi^) compared to ipsilateral CA3 (CA3^ipsi^) and contralateral CA1 (CA1^contra^) (scale bars: 50 µm). **(D)** Representative 2-photon micrographs showing that microglia at the CA1^ipsi^ injection site display a more amoeboid morphology (left) than microglia in CA3^ipsi^ (middle) and CA1^contra^ (right) regions (scale bars: 10 µm). **(E)** Input-output curves showing that 470 nm light stimulation evoked field excitatory postsynaptic potentials (fEPSPs) in CA3^ipsi^ *stratum radiatum*, but not in CA1^contra^ *stratum radiatum.* Example traces are displayed for 2 mW light stimulation (scale bar: 0.5 mV, 5 ms).

**Figure S4.**
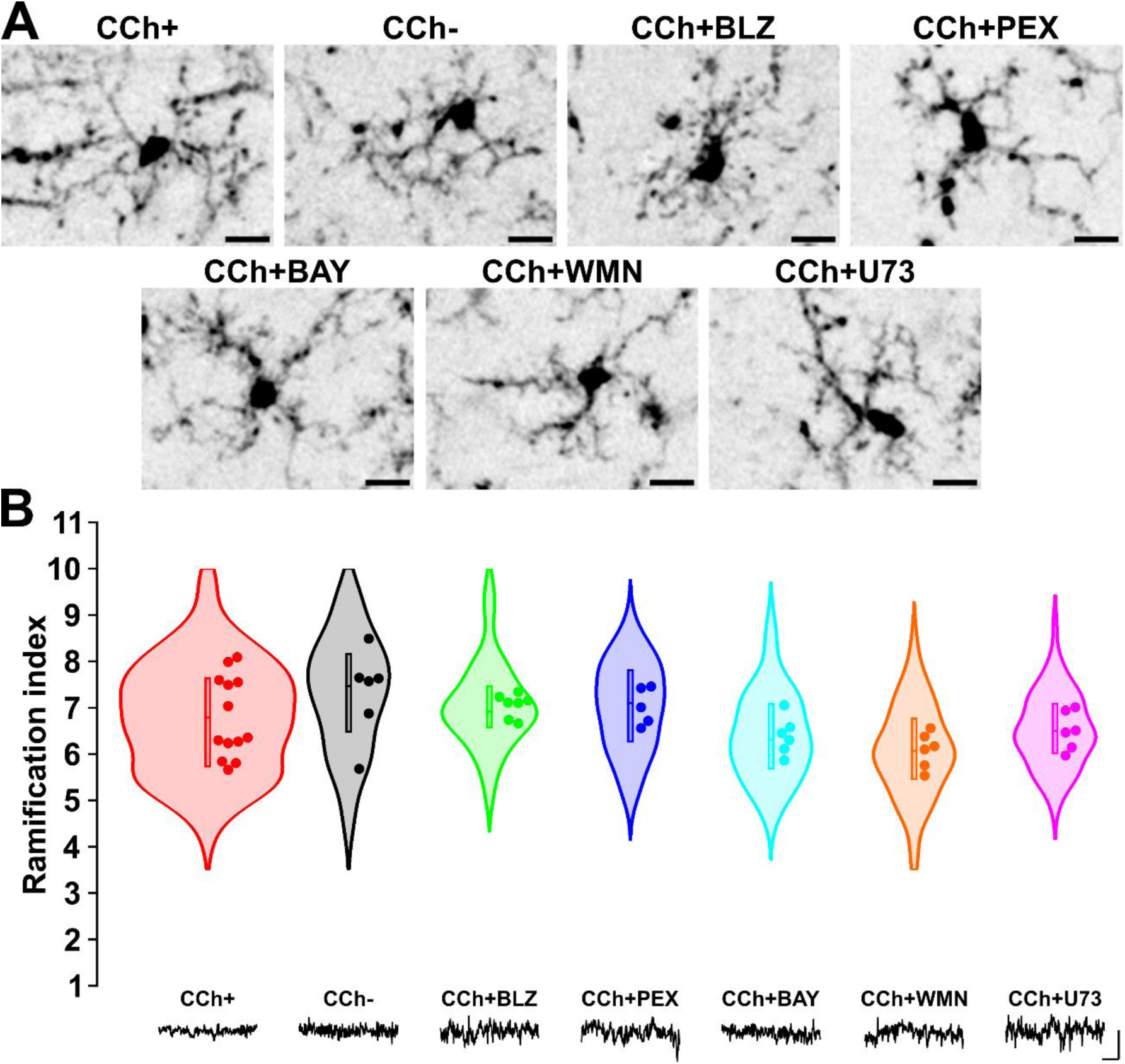
Microglial morphology is unaltered by BLZ-945, pexidartinib, bay 11-7082, wortmannin, or U73122. **(A)** 2-photon micrographs of example microglia PRh of CX3CR1^+/GFP^ slices following 30 min incubation with carbachol (CCh+), or co-application of CCh with BLZ-945 (CCh+BLZ), pexidartinib (CCh+PEX), bay 11-7082 (CCh+BAY), wortmannin (CCh+WMN), or U73122 (CCh+U73). Microglia morphology in vehicle slices without drug application (CCh-) is also shown (scale bars: 10 µm). **(B)** PRh microglia RIs under each experimental condition. Traces illustrate example LFPs per condition (scale bar: 0.1 mV, 50 ms). See **Table S3** for statistical analysis. See **Table S4** for working drug concentrations.

**Table S1.**
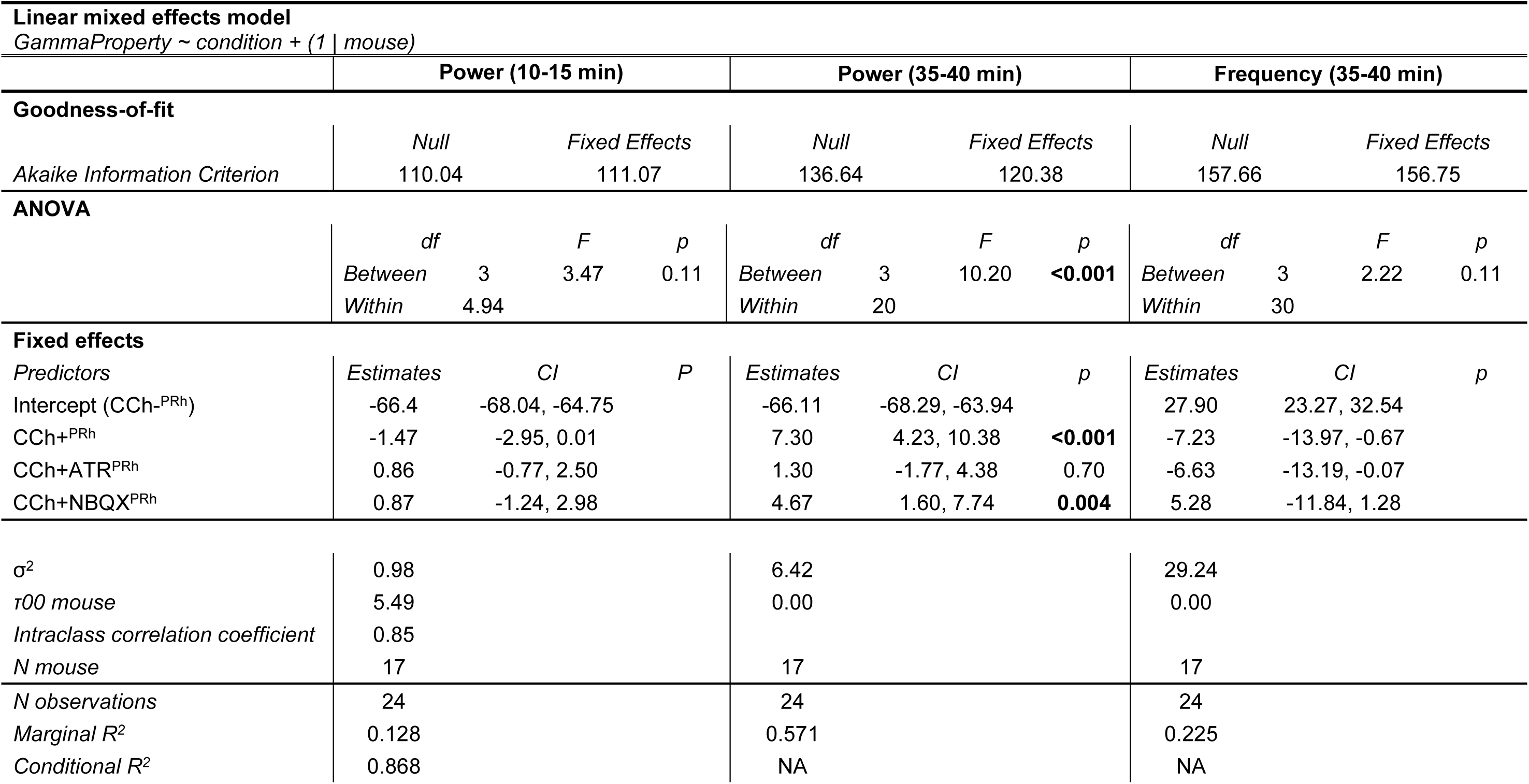
Linear mixed effects models comparing baseline gamma power (10-15 min) and CCh evoked gamma power and frequency (35-40 min) between vehicle control (CCh-), CCh treatment (CCh+), CCh+ATR treatment, and CCh+NBQX treatment conditions in the perirhinal cortex (PRh) of CX3CR1^+/GFP^ mouse cortico-hippocampal slices. To evaluate model goodness-of-fit, the table displays Akaike Information Criterion values for null and mixed effects (condition) models. Main fixed effects were evaluated by one-way ANOVA. The table displays the degrees of freedom (*df*), test statistic (*F*), and significance (*p*) values. For fixed effects, the table displays the estimates, confidence intervals (CI) and *p* values for *post hoc* tests (Dunnett’s comparison to CCh-^PRh^ with Holm-Bonferroni correction). For random effects, the table displays residual variance (σ2), mouse variance (τ00), and the intraclass correlation coefficient (ICC). Marginal R^2^ refers to the variance explained by fixed effects only. Conditional R^2^ refers to variance explained by fixed and random effects combined.

**Table S2.** Linear mixed effects model comparing microglia ramification indices (RI) between vehicle control (CCh-), CCh treatment (CCh+), CCh plus atropine treatment (CCh+ATR), and CCh plus NBQX treatment (CCh+NBQX) conditions in the perirhinal cortex (PRh) of CX3CR1^+/GFP^ mouse cortico-hippocampal slices. To evaluate model goodness-of-fit, the table displays Akaike Information Criterion values for null and mixed effects (condition) models. Main fixed effects were evaluated by one-way ANOVA. The table displays the degrees of freedom (*df*), test statistic (*F*), and significance (*p*) values. For fixed effects, the table displays the estimates and confidence intervals (CI). For random effects, the table displays residual variance (σ2), slice variance (τ00 slice), and the intraclass correlation coefficient (ICC). Marginal R^2^ refers to the variance explained by fixed effects only. Conditional R^2^ refers to variance explained by fixed and random effects combined.

| Linear mixed effects model |  |  |  |
| --- | --- | --- | --- |
| RI ~ condition + (1 \| slice) |  |  |  |
| Goodness-of-fit |  |  |  |
|  | Null | Condition |  |
| Akaike information criterion | 670.38 | 674.89 |  |
| ANOVA |  |  |  |
|  | df | F | p |
| Between | 3 | 0.428 | 0.74 |
| Within | 20 |  |  |
| Fixed effects |  |  |  |
| Predictors | Estimates | CI |  |
| Intercept (CCh-<sup>PRh</sup>) | 6.21 | 5.82, 6.60 |  |
| CCh+<sup>PRh</sup> | 0.25 | -0.29, 0.80 |  |
| CCh+ATR<sup>PRh</sup> | 0.07 | -0.48, 0.61 |  |
| CCh+NBQX<sup>PRh</sup> | -0.03 | -0.58, 0.52 |  |
| Random effects |  |  |  |
| σ<sup>2</sup> | 0.85 |  |  |
| τ00 slice | 0.15 |  |  |
| Intraclass correlation coefficient | 0.15 |  |  |
| N slice | 24 |  |  |
| N observations | 240 |  |  |
| Marginal R<sup>2</sup> | 0.012 |  |  |
| Conditional R<sup>2</sup> | 0.158 |  |  |

**Table S3.**
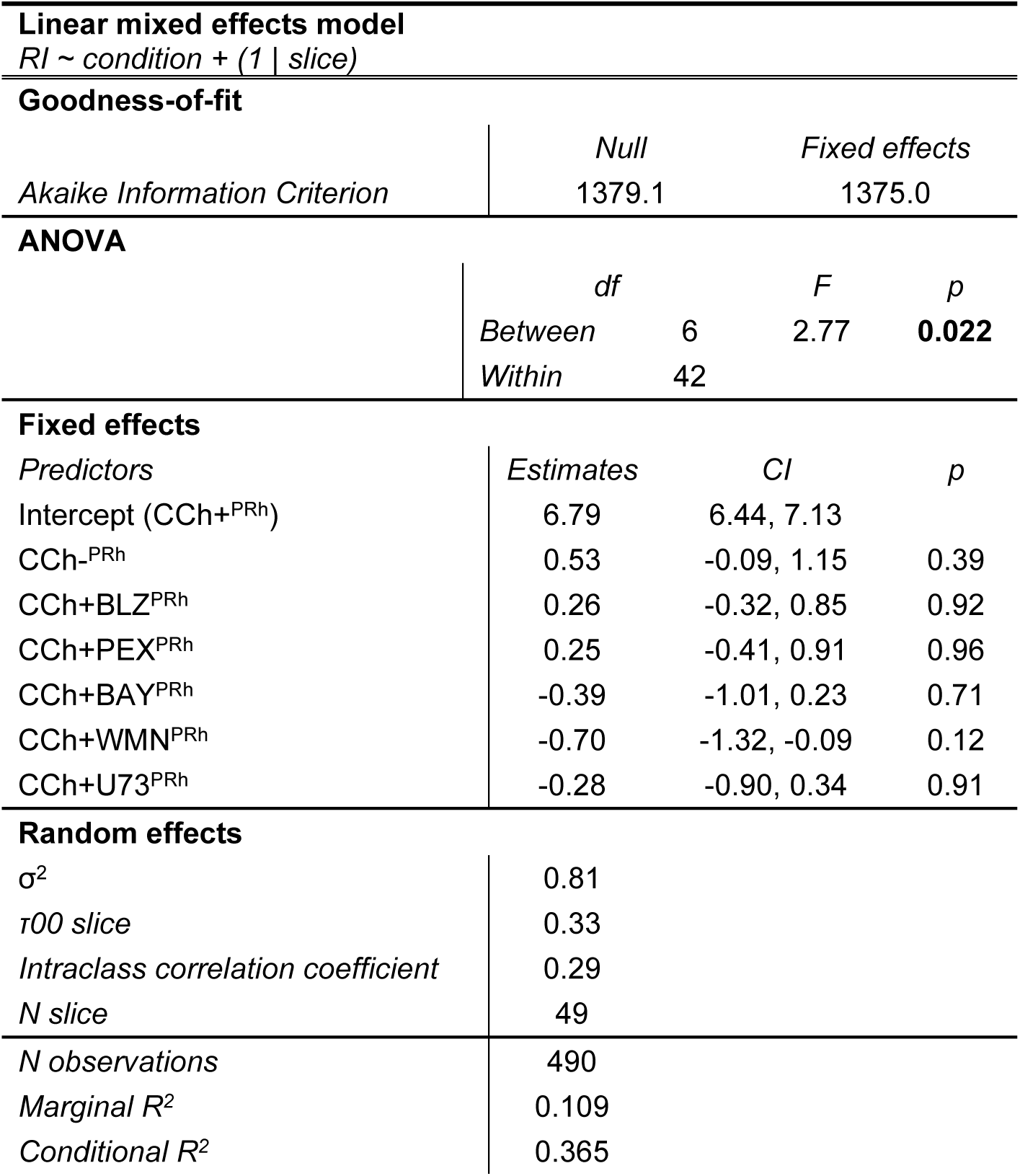
Linear mixed effects model comparing microglia ramification indices (RI) between CCh treatment (CCh+), vehicle control (CCh-), CCh plus BLZ-945 treatment (CCh+BLZ), CCh plus pexidartinib treatment (CCh+PEX), CCh plus bay 11-7082 treatment (CCh+BAY), CCh plus wortmannin treatment (CCh+WMN), and CCh plus U73122 treatment (CCh+U73) conditions in the perirhinal cortex (PRh) of CX3CR1^+/GFP^ mouse cortico-hippocampal slices. To evaluate model goodness-of-fit, the table displays Akaike Information Criterion values for null and mixed effects (condition) models. Main fixed effects were evaluated by one-way ANOVA. The table displays the degrees of freedom (*df*), test statistic (*F*), and significance (*p*) values. For fixed effects, the table displays the estimates, confidence intervals (CI), and *p* values for *post hoc* tests (Dunnett’s comparison to CCh+ with Holm-Bonferroni correction). For random effects, the table displays residual variance (σ2), slice variance (τ00 slice), and the intraclass correlation coefficient (ICC). Marginal R^2^ refers to the variance explained by fixed effects only. Conditional R^2^ refers to variance explained by fixed and random effects combined.

**Table S4.** Compounds used in pharmacology experiments.

| Compound name | Biological activity | Concentration (μM) | Supplier | Product code |
| --- | --- | --- | --- | --- |
| Atropine | Muscarinic receptor antagonist | 10 | Merck Life Sciences | A0132 |
| Bay 11-7082 | IκB inhibitor | 100 | Merck Life Sciences | B5556 |
| BLZ-945 | CSF-1 receptor antagonist | 0.5 | HelloBio | HB4634 |
| Carbachol | Cholinergic receptor agonist | 20 | Tocris Bioscience | 2810 |
| NBQX | AMPA receptor antagonist | 10 | HelloBio | HB0443 |
| Pexidartinib | CSF-1 receptor antagonist | 1 | Tocris Bioscience | 7590 |
| U73122 | Phospholipase C inhibitor | 10 | Tocris Bioscience | 1268 |
| Wortmannin | PI3 kinase inhibitor | 10 | Tocris Bioscience | 1232 |

